# The tumor suppressor p53 mutational status controls epithelial 3D cell growth under mechanical compression

**DOI:** 10.1101/2025.10.23.681444

**Authors:** Mickael Di-Luoffo, Silvia Arcucci, Nicole Therville, Tristan Marty, Romina D’Angelo, Maria Chaouki, Benoît Thibault, Pascal Swider, Pauline Assemat, Morgan Delarue, Julie Guillermet-Guibert

## Abstract

**Context:** Solid tumors are subjected to mechanical stimuli arising from their growth in confined environments. Growth-induced pressure builds up in tumors such as pancreatic cancer and rises alongside the occurrence of genetic alterations during tumorigenesis. This study aims to understand the so far unknown relationship between genetic alterations and cancer cell behavior under compressive stress.

**Results:** Using isogenic cell lines with engineered p53 mutations, we showed that the p53 background influences cell response to compression. Tumor growth under compression increased in cells harboring a mutated-truncated p53 form. This mutation blocked caspase 3 cleavage and promoted survival and growth through PI3K-AKT activation and dysregulation of c-FOS and FOSB transcription factors network. Mutated-truncated p53 cells displayed a unique behavior and heightened an activation state under compression.

**Conclusion:** Mechanical compression and p53 mutations together drive tumor growth. p53 status could be a biomarker for predicting tumor adaptation to mechanical stress and efficiency of therapies targeting mechanosensitive pathways.

**Teaser:** Mechanical compression and p53 mutations together enhance cancer cell survival and growth, driving solid tumor progression.

## Introduction

Mechanical stresses and strains are inherent to physiological tissues and largely associated to solid cancers. At a cellular scale, all cells are continuously exposed to mechanical stimuli. Mechanical stresses can be transmitted to internal structures, including the nucleus (*1*). There are three types of mechanical stresses: shear, tensile and compressive stresses (*2*). These stresses originate from cell-cell interactions, from cell-extracellular matrix (ECM) interactions and eventually from interstitial flows (*3*). During solid cancer development, the rapid growth and proliferation of tumor cells associated with accumulation of ECM in a tissue (*4*) increases cell confinement leading to a growth-induced pressure, which in turn increases the compressive stress applied to tumor and stromal cells (*5*). When mechanical stress is applied to a cell, it triggers a mechanotransduction response that leads to alterations in gene expression, mutations and/or pathways dysregulation. Under physiological conditions, epithelial cells respond to compressive strains and stresses by limiting their proliferation, nevertheless such stimuli are relatively low in homeostasis conditions (*6*). In tumors such as pancreatic cancer (PDAC), the cells grow despite the build-up of compressive stress (*4*). Confinement-induced compression also increases resistance to chemotherapy, partly because it sustains cell survival and reduces proliferation, thus decreasing the effectiveness of cytotoxic and cytostatic drugs (*7, 8*).

Increase of mechanical stimuli and occurrence of genetic alterations are concomitant in solid cancers development (*9, 10*). One of the mutations associated with solid cancers is the mutation of the small GTPase KRAS associated to MAP Kinase (MAPK) and phosphoinositide 3-kinase (PI3K) pathway. PI3K pathway is altered in more than 50% of all cancers (*11*). In physiological conditions, KRAS switches between inactivated GDP-bound form and an active form bound to GTP (*12*). The KRAS-GTP active form directly interacts with downstream effectors in MAPK pathway and different PI3K isoforms (*13–15*). In most cancers, oncogenic KRAS genetic alterations are generally occurring in the initial step of solid cancer progression (*16, 17*). Solid cancers harboring the highest mutation rates on *KRAS* are pancreatic cancer (pancreatic ductal adenocarcinoma, PDAC), found in about 90-95% of cases (*12*), colorectal cancer (CRC) in 35-45% of tumors (*13*) and lung adenocarcinoma in 20-30% of cases (*18*).

Associated with *KRAS* mutations, mutations in the *TP53* gene occur in more than 50% in head and neck, lung, colorectal, ovarian, esophageal and pancreatic cancers (*19–23*). In the case of PDAC, mutations in the *TP53* gene are found in more than 60% of patients (*24*). One of the most common mutations found in humans is p53^R175H^ mutation (*25*), along with its mouse homologue p53^R172H^ (*24, 25*). This alteration leads to a loss of function of p53 as a genome guardian associated with a gain of function of the protein, conferring proliferative, invasive, and migratory capabilities to cancer cells, as well as resistance to chemotherapeutic agents (*26*). PDAC is one of the cancers with the most frequent genetic alterations of p53, and some of these mutations or insertions/deletions lead to a translational frameshift inducing insertion of a STOP codon in its sequence and the translation of a truncated p53 protein (*27, 28*).

Over the last decades, researchers have been interested in the effect of mechanical stresses and strains during in both physiological and pathological conditions (*29–31*). In particular, deciphering the effect of mechanical stresses that emerge during solid cancer development has become increasingly timely (*32, 33*). However, the causal association between the combined genetic and mechanical context and solid cancer progression is poorly studied. Here, we aim at understanding the importance of various genetic context (such as *KRAS* and *TP53* mutations) on tumor behavior under compressive stimuli, using PDAC as a paradigmatic model of heightened confinement.

The genetic context is well established in the diagnosis and care of most patients with solid cancers: using Next Generation Sequencing or DNA Methylation Analysis, some cancer treatments are based on identified genetic alterations (*34, 35*). Our work identified selective mutants of p53 that sustain growth under compression and favors oncogenic PI3K activation. This knowledge will help to move towards a personalized patient care, taking into account both genetics and mechanics for patient-personalized cancer therapy.

## Results

### 1. p53 mutation prevents the 3D growth reduction normally induced by confinement in PDAC spheroids

To answer our question, we positioned our experimental setup to mimic solid cancer development (Figure 1A). While the genetic alterations and genome instability prompt the evolution of lesions, the induced mechanical context leads to the confinement of growing cancer cells. A paradigmatic model to study this stage transition is pancreatic cancer (PDAC) (*25, 26*). To model this, we developed isogenic cell lines from parental *KRAS^G12D^* murine PDAC cells, stably transfected with a doxycycline-inducible Tet-ON system. We prompted a stable expression of GFP in cells called K^+^ (*KRAS^G12D^*) combined or not with an enhanced expression of a mutated form of p53 (p53^R172H^) in cells called K^+^p53^M^ (Figure 1B; Supp Figure 1A). DNA sequencing blasted to p53^WT^ sequence validated that PDAC cells were transfected with empty plasmid or plasmid containing p53^R172H^ mutation (Supp Figure 1B). Further, p53 level in K^+^p53^M^ cells was 3.3-fold increased compared to isogenic empty vector clones (K^+^) (Supp Figure 1C).

**Figure 1.**
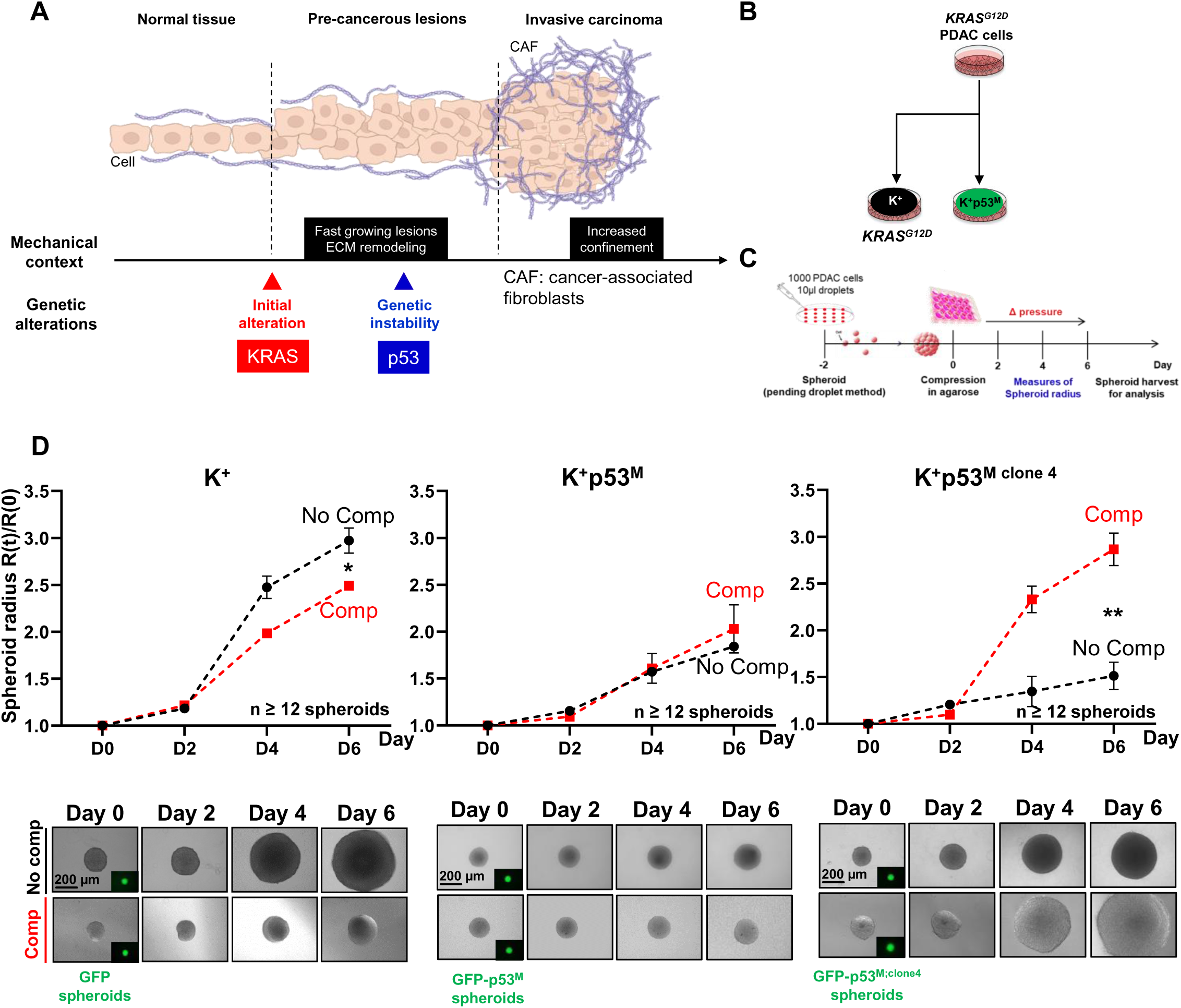
p53 mutation prevents the 3D growth reduction normally induced by confinement in PDAC spheroids. **A.** Current model of solid cancer development: normal cells evolve to pre-cancerous lesion, then to invasive carcinoma; the mechanical context is deeply modified in fastly growing lesions, as extracellular matrix (ECM) remodels during those stage progressions. Stiffening and swelling of the matrix lead to the confinement of cancer initiating and cancer cells. The genetic context, such as KRAS and p53 genetic alterations, evolves during carcinogenesis and influences tumor development. CAF: cancer associated fibroblast. **B.** Schematic of isogenic cell line creation: from parental KRAS^G12D^ pancreatic cancer cells (PDAC cells), the Tet-ON stable expression system allowed protein expression from empty vector (GFP only in mutated KRAS^G12D^ background: K^+^) or from p53^R172H^ expression vector (KRAS^G12D^; p53^R172H^: K^+^p53^M^). **C.** Schematic of experimental setup: 1000 cells were placed on the cover of 100 mm petri dish. The cover was flipped over and using the pending droplet method, spheroids were formed in the droplet meniscus during 2 days. After spheroids generation, they were confined into 1% low gelling temperature agarose at day 0 and for 6 days in order to apply on spheroids a growth induced pressure (Δ pressure). During the 6 days, pictures were taken and spheroids radius were measured. Spheroids were harvested at day 6 for future analysis. **D.** K^+^ genotype spheroids (left panel), three clones of K^+^p53^M^ genotype spheroids (center panel) and one clone of K^+^p53^M^ genotype spheroids (right panel) were confined inducing compressive stress (Comp) compared to not compressed (No Comp). The spheroids radius R(t)/R(0) were analyzed for 6 days with Day 0 corresponding to confinement initiation. The growth curves are presented depending on the 3D growth behavior of spheroids under confinement: confinement-induced significantly decreased spheroids growth (corresponding to K^+^ genotype, left panel); no change in growth (3 out of 4 clones from K^+^p53^M^ genotype, center panel); confinement-induced significantly increased 3D growth (1 out of 4 clones from K^+^p53^M^ genotype, right panel). GFP expression related to empty vector or p53^M^ expressions in spheroids from day 0 to day 6. Scale bar corresponds to 200µm. Results were represented as mean, +/- SEM, n≥12 spheroids from independent cultures per condition. *p-value<0.05; **p-value<0.01. One representative spheroid picture was shown in each condition.

Four clones of PDAC murine cells harboring *KRAS^G12D^* single mutation (K^+^) or harboring *KRAS^G12D^; p53^R172H^*mutations (K^+^p53^M^) were used to test the effect of 3D confinement (Figure 1C). After 2 days, K^+^ or K^+^p53^M^ spheroids made with the pending droplet method were deposited onto a 1% low gelling temperature agarose cushion to allow their free growth; others were embedded into a 1% low gelling temperature agarose (Figure 1C). Both media were supplemented with doxycycline for 2 days. Confined growth in agarose induced a growth-induced pressure as estimated in Supp Figure 1D (detailed in M&M §*Spheroids generation and agarose confinement* (*8*)), generating mechanical compressive stress on cells. By following the evolution of spheroids over 6 days, we first identified the effect of such stress on long-term 3D growth. GFP expression was stable in all spheroids (Figure 1D, see inset pictures). Free-growing K^+^p53^M^ spheroids grew slower over time than K^+^ spheroids (No Comp vs No comp in Figure 1D left and center panel, graphs and pictures). However, the 3D confinement significantly reduced the 6 day-spheroid radial growth by 16% in K^+^ genotype (Comp vs No comp in Figure 1D left panel, graphs and pictures) and did not affect spheroid growth in 3 out of 4 clones of K^+^p53^M^ genotype (Figure 1D center panel, graphs and pictures). The confined K^+^p53^M^ spheroids showed a consistent 3D growth advantage at long term compared to their free growing counterparts (calculated growth rates are shown in Supp Figure 1E). In all genotypes, the early growth-induced pressure can be estimated in the early time points after careful rheological measurement of agarose properties and finite-element simulations of a spheroid growth in such material (*8*). Mechanical compressive stress was in the order of kPa, 1.6kPa in K^+^ genotype and 0.6 kPa in K^+^p53^M^ genotype (Supp Figure 1D). Surprisingly, 1 clone (K^+^p53^M;clone^ ^4^) saw its long-term radial growth significantly increased under confinement (+60% in K^+^p53^M;clone^ ^4^ compressed spheroids compared to uncompressed) (Figure 1D right panel, graphs and pictures).

The K^+^ clones are sensitive to confinement, and their long-term confined 3D growth is reduced, while three K^+^p53^M^ clones are insensitive to confinement and one K^+^p53^M^ clone increased its 3D growth under confinement. The embedding in the rigid and inert matrix is providing a proliferative advantage to these spheroids.

### 2. The heightened growth under confinement is explained by a truncation of p53 protein

We observed that K^+^p53^M;clone^ ^4^ grew faster in agarose (Figure 1D). The western blot analysis of p53 protein revealed that the protein level of the full-length form of p53 was equally increased in the four clones (K^+^p53^M^) compared to the K^+^ clone (Figure 2A upper bands). A lower band was present at approximately 28 kDa size, which could correspond to a truncated form of p53. This truncation of p53 is found endogenously in humans and named p53^psi^ form (*27*). This band was barely present in the K^+^ clone and in the first three K^+^p53^M^ clones. However, its level was increased in the fourth clone (K^+^p53^M;clone^ ^4^) (Figure 2A lower bands). In the genomics data of 20 cancer types from TCGA cohorts (between 185 and 1098 cases per cancer), genetic alterations encoding for truncated forms of p53 were present in 16.5% to 2% of cases depending on the cancer observed (from esophageal squamous cell cancer to myeloma) (Figure 2B). In TCGA-PDAC cohort (PAAD-TCGA), a genetic alteration encoding for p53 truncation was found in 11.2% of cases, which places PDAC at the 6^th^ rank of the most truncated cancers for p53 (Figure 2B). Among the 145 PDAC patients from the TCGA, 70% harbored a p53 mutation. Among the 70% of mutated p53 in PDAC cohort, 50% were missenses mutations, 12% truncations in other exons and 8% truncations in exon 6 of p53 (Figure 2C).

**Figure 2.**
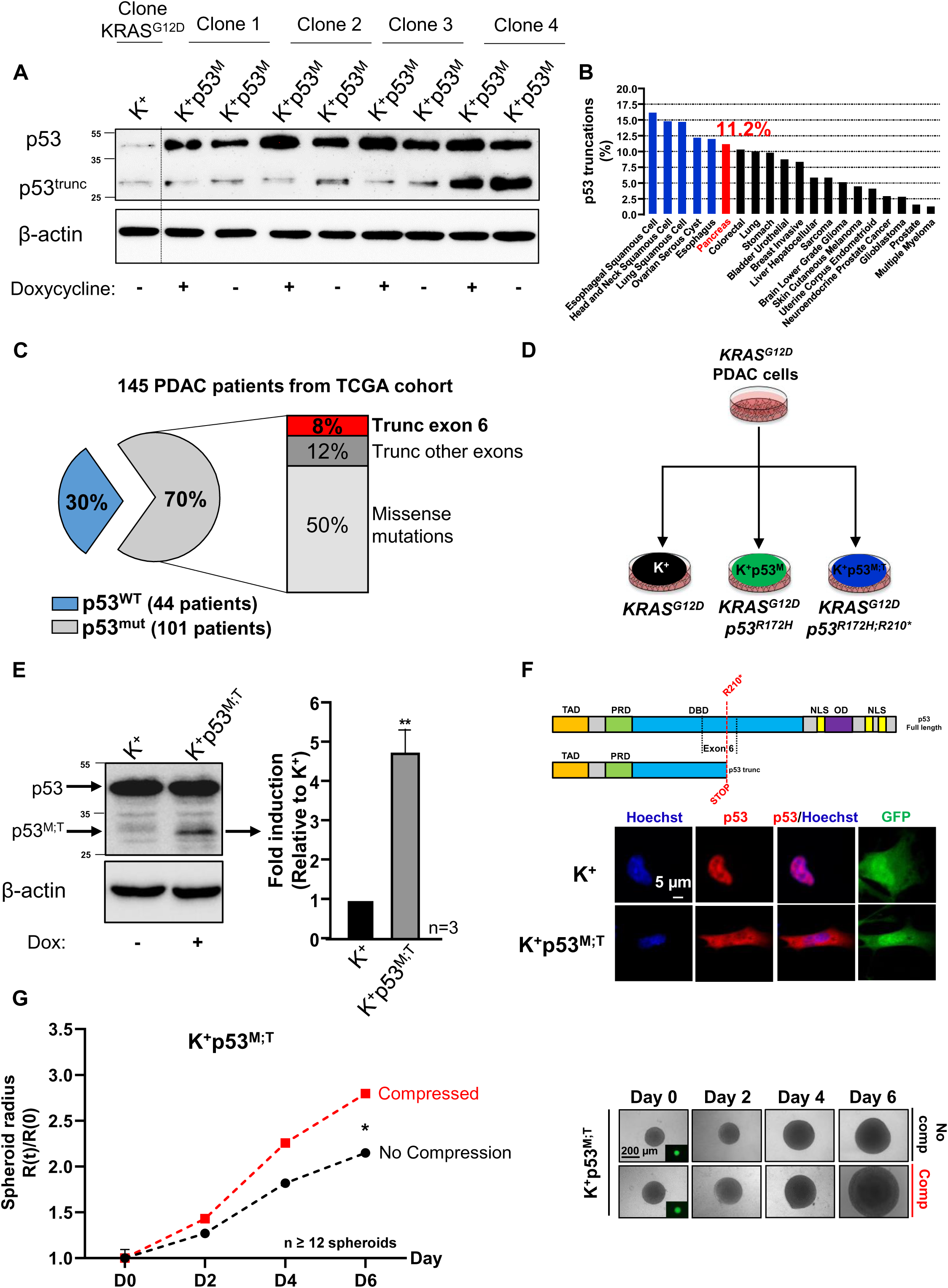
The heightened growth under confinement is explained by a truncation of p53 protein. **A.** Representative western blots of p53 and endogenous p53 truncated form (p53^trunc^) in KRAS^G12D^ (K^+^) (first lane) and KRAS^G12D^; p53^R172H^ (K^+^p53^M^) clone 1,2,3 and 4 (lanes 2 to 8). β-ACTIN was used as loading control. Doxycycline (500ng/mL) was used to induce Tet-ON stable expression of p53^R172H^ in K^+^; p53^M^ clones. **B.** Gene alterations leading to expression of truncated forms of p53 are found in 20 cancers in the TCGA cohort, including pancreatic cancer (6th position in 20 cancer types from TCGA cohort). In addition, pancreatic cancer is the first most frequent cancer in which a mutation encoding a truncation of p53 in exon 6 is found among the 20 cancers in the TCGA cohort. **C.** In 145 patients with pancreatic cancer (TCGA cohort), 20% of p53 mutations result in a truncation of the protein and 8% are specific to exon 6. **D.** From KRAS^G12D^ pancreatic cancer cells (PDAC cells), the Tet-ON stable expression system allows protein expression of GFP in empty vector (with mutated KRAS^G12D^ background: K^+^) or of p53^R172H^ and GFP expression (in KRAS^G12D^backgound p53^R172H^: K^+^p53^M^) or of p53^R172H;R210*^ and GFP expression (KRAS^G12D^ in background with p53^R172H;;R210*^: K^+^p53^M;T^) . **E.** Representative western blots of p53 and genetically engineered p53 truncated form (p53^M;T^) in KRAS^G12D^ background (K+) (first lanes) and KRAS^G12D^; p53^R172H;R210*^ (K^+^; p53^M;T^) clone (lane 2). β-ACTIN was used as loading control. Doxycycline (Dox) (500ng/mL) was used to induced Tet-ON stable expression of p53^R172H;R210*^ in K^+^; p53^M;T^ clone. p53^M;T^/β-ACTIN quantitative analyses were performed using ImageJ software. Results are presented as mean, +/- SEM, n≥3. **p-value<0.01. **F.** The truncation in R210* position in exon 6 is predicted to remove part of the DNA-binding domain (DBD), the two nuclear localization domains (NLS) and the oligomerization domain (OD) of p53 protein (up panel). After immunofluorescence anti-p53, the genetically engineered truncated form of p53 (K+; p53^M;T^) was found in the cytosol while the p53 WT form was found in the nucleus of pancreatic (K^+^) cells. Nuclear signal corresponds to Hoechst and GFP is expressed in the cytosol of the transfected pancreatic cells. **G.** K^+^; p53^M;T^ genotype spheroids were confined inducing compressive stress (Compressed) compared to no compression condition (No Compression). The spheroids radiuses were measured at days 0, 2, 4, and 6 and day 0 (D0) corresponding to initiation of compression. This confinement significantly increased spheroids growth (dotted red curve (Compressed) compared to black dotted curve (No Compression)) (left graph). Representative images of K^+^; p53^M;T^ not compressed (No Comp) (high right panel) and compressed (Comp) (low right panel) spheroids were shown. Scale bar corresponds to 200µm. GFP expression in spheroids from day 0 to day 6 was shown. Results are presented as mean, +/- SEM, n≥12 spheroids per condition. *p-value<0.05.

We hypothesized that the increased level of truncated form of p53 found in the (K^+^p53^M;clone^ ^4^) could be responsible for the increased 3D growth under confinement. We next developed another isogenic cell line from the parental PDAC murine cells harboring *KRAS^G12D^* single mutation with a stable expression of a mutated-truncated form of p53 in exon 6 (p53^R172H;R210*^, called K^+^p53^M;T^) using the doxycycline-inducible Tet-ON system (Figure 2D). Western blot analysis showed that the truncated form of p53 was significantly and 4.6-fold higher in the mutated/truncated p53 cells (K^+^p53^M;T^) compared to the KRAS^G12D^ (K^+^) mutated cells (Figure 2E). A truncation by insertion of STOP codon in exon 6, invalidates the DNA binding domain and removes the nuclear localization domains (Figure 2F), which could amplify the gain-of-function of missense mutants (*27, 28*). In accordance, we demonstrated that p53 truncated in exon 6 (K^+^p53^M;T^) presented a localization restricted to cytosol, while wild-type p53 was mainly found in the nucleus (*27*) (Figure 2F).

In addition, confined K^+^p53^M;T^ spheroid radial growth was significantly increased by 28% (Comp) compared to free-growing K^+^p53^M;T^ spheroids (Figure 2G); in this context, confined 3D growth allowed the accumulation of a growth-induced pressure of 3.9 kPa (Supp Figure 1D). We tested whether the 3D growth advantage provided by confinement in p53 mutant cells was due to an intrinsic bias caused by genotype. In 2D , K^+^p53^M^ cell numbers increased significantly and more rapidly than the K^+^ cells (Supp Figure 1F). The K^+^p53^M;T^ and K^+^p53^M^ ^clone4^ cells, however, had a slightly reduced curve compared to the control K^+^ cells, this delay was also observed in free spheroid growth (Supp Figure 1F, Figure 1D, Figure 2G).

The increased 3D growth under confinement in a rigid matrix of clone 4 is phenocopied by a truncation of p53 associated with a p53 mutation. Intrinsic 2D growth rate of K^+^p53^M;T^ and K^+^p53^M^ ^clone4^ cells did not explain the observed increased 3D growth rate under confinement. We next aimed to understand more precisely the mechanisms associated with 3D growth increase under confinement by mutated-truncated p53.

### 3. Mutated-truncated p53 cells lose the growth-inhibitory early AP-1 pathway gene response under confinement

In order to better understand the intracellular mechanisms associated with the 3D growth advantage under confinement, we investigated the early gene response. Spheroids of either K^+^, K^+^p53^M^ or K^+^p53^M;T^ cells were confined and analyzed by RNA sequencing at an early time point (D2), prior to observed differences in long-term 3D growth (Figure 3A). The principal component analysis (PCA mapping) allowed to identify two groups of samples: the group containing K^+^ and K^+^p53^M^ spheroids and another group including K^+^p53^M;T^ spheroids (Figure 3A, Supp Figure 2A). A difference between confined and unconfined conditions was also clearly identified in K^+^p53^M;T^ spheroids (Figure 3A). We focused our analysis on the significantly upregulated genes that were less prone to bias in sample preparation. The expression of the top 15 genes upregulated by compressive stress in K^+^, K^+^p53^M^ spheroids differed largely between each genotypes (Figure 3B, Supporting Table 1). In K^+^p53^M;T^ genotype, we identified, amongst the top 15 genes, two genes of AP-1 family (Activator Protein 1), *c-Fos* and *FosB* (FBJ Murine Osteosarcoma Viral Oncogene Homolog), which exhibited higher expression under confinement compared to K^+^ and K^+^p53^M^ genotypes (Figure 3B, 3C). We next analyzed relative mRNA levels by RT-qPCR. *c-Fos* and *FosB* mRNA expressions were significantly increased by 5.4 and 2.2-fold in K^+^p53^M;T^ spheroids under confinement (Figure 3D); however, *c-Fos* was also found increased by 3.7 and 3.4 in K+ and K^+^p53^M^ cells, respectively (Figure 3D). Those RT-qPCR results are in line with the global RNAseq analysis, in which we observed that the top regulated genes in K^+^p53^M;T^ spheroids were also globally upregulated to the same scale in K^+^ and K^+^p53^M^ (Figure 3B,C).

**Figure 3.**
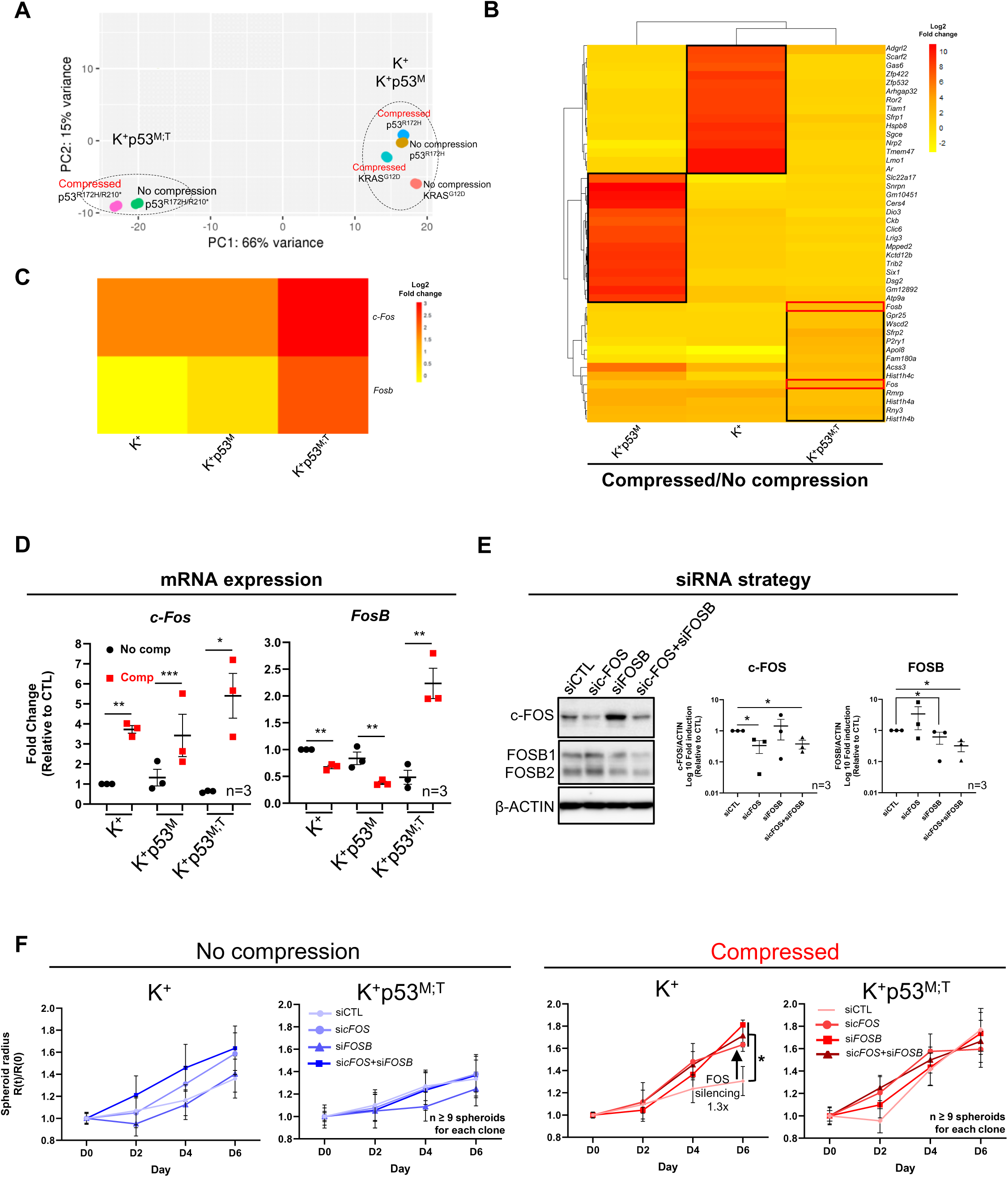
Mutated-truncated p53 cells lose the growth-inhibitory early AP-1 pathway gene response under confinement. **A.** Principal Component Analysis (PCA) mapping represents the distribution of RNA sequencing samples divided into two groups: on one hand KRAS^G12D^ and KRAS^G12D^; p53^R172H^ (K^+^p53^M^), compressed and not compressed conditions and on the other hand the KRAS^G12D^, p53^R172H;R210*^ (p53^M;T^) compressed and not compressed conditions (day2). n=3 pools of 24 spheroids per condition. **B.** Heatmap representation of top 10 overexpressed genes in compressed compared to not compressed conditions in K^+^, K^+^p53^M^ and K^+^p53^M;T^ genotypes. Black squares represent top overexpressed genes in each genotype and red squares represent *c-Fos* and *Fosb* overexpression in K^+^p53^M;T^ genotype **C.** Heatmap representation of *c-Fos* and *Fosb* gene expression in K^+^, K^+^p53^M^ and K^+^p53^M;T^ genotypes. Gene expression was represented as log2 Fold Change values in compressed compared to not compressed conditions. NA was represented as log2FC=0. **D.** mRNA expression of *c-Fos* and *Fosb* in compressed (Comp, red squares) *vs* not compressed (No comp, black dots) in K^+^, K^+^p53^M^ and K^+^p53^M;T^ genotypes. Results are presented as mean, +/- SEM, n=3 x24 spheroids per condition. *p-value<0.05; **p-value<0.01; ***p-value<0.001. **E.** Representative western blots of c-FOS and FOSB protein expression after RNA silencing against c-FOS (sicFOS), against FOSB (siFOSB) and a combination of siRNA against c-FOS and FOSB (sicFOS+siFOSB). SiCTL corresponds to smart pool of siRNA scrambled. β-ACTIN was used as loading control. Quantitative analyses were performed using ImageJ software and represents the fold induction of c-FOS/ β-ACTIN and FOSB/ β-ACTIN. Results are presented as mean, +/- SEM, n=3 per condition. *p-value<0.05. **F.** K^+^ K^+^p53^M;T^ genotype spheroids were treated using silencing RNA against c-FOS (sicFOS), against FOSB (siFOSB) and a combination of siRNA against c-FOS and FOSB (sicFOS+siFOSB), confined (Compressed) compared to no compression condition (No Compression). SiCTL corresponds to smart pool of siRNA scrambled. The spheroids radiuses were measured for 6 days and day 0 (D0) corresponding to initiation of compression. Results were represented as mean, +/- SEM, n≥12 spheroids per condition. *p-value<0.05.

To demonstrate the role of these two transcription factors in 3D growth under confinement, we developed a c-FOS and FOSB knock down strategy (alone or in combination), using siRNAs directed against each of them (Figure 3E). To validate the knock down strategy, we performed a western blot against c-FOS and FOSB from transfected cells with scramble siRNA or siRNA against c-FOS, FOSB and combination of both. SiRNA against c-FOS and combination of siRNA against c-FOS and FOSB significantly decreased c-FOS protein expression (Figure 3E). Moreover, siRNA against FOSB and combination of siRNA against FOSB and c-FOS significantly decreased FOSB protein level compared to scramble siRNA (siCTL) (Figure 3E).

In K^+^ and K^+^p53^M;T^ spheroids, siRNA against c-FOS and FOSB or combination of both did not affect spheroids growth in free growth condition (Figure 3F left panels). However, FOS factors silencing (c-FOS, FOSB alone and both) significantly increased spheroid growth by 1.3 fold in confined K^+^ spheroids but not in confined K^+^p53^M;T^ spheroids (Figure 3F right panels). The compressed FOS-silenced K^+^ spheroid growth behaved as compressed K^+^p53^M;T^ spheroids and free-growing K^+^ spheroids (Figure 3F).

We here identified a mechanism responsible for reduced 3D growth in confined K^+^ spheroids which is abrogated by expression of p53^M;T^, while the additional overexpression of both c-FOS, FOSB in confined K^+^p53^M;T^ spheroids could be a compensatory response to the absence of FOS factor cellular effects.

### 4. Confinement activates differently PI3K-AKT pathway in mutated-truncated p53 cells

We next hypothesized that, in K^+^p53^M;T^ genotype, a signaling adaptive response could be responsible for the increased growth fitness in response to confinement. We thus analyzed in an unbiased manner the gene signatures enriched in compressed condition. While all genotype harbored enrichment in signatures associated to DNA regulation (GO terms, Supp Figure 2B) further validating our findings in Figure 3, we also observed a strong enrichment of signatures linked to signalling in K^+^p53^M;T^ spheroids under confinement *vs* free-growth (Figure 4B). We identified an enrichment in genes controlling cell adhesion *via* the plasma membrane and pathways associated with tyrosine kinase receptors (TKR) that are partly regulated or translated into biochemical signals by PI3K pathway and MAPK (KRAS) pathway, while genes controlling angiogenesis and cell migration pathways were enriched in a lower level (Figure 4B). As a major pathway downstream TKR and cell adhesion molecules (*36*), the expression of genes in the REACTOME_PI3K-AKT signature in cancer was analyzed. Under confinement, the genetic background of spheroids (K^+^, K^+^p53^M^ or K^+^p53^M;T^) modulated the quantitative and qualitative gene expression in the PI3K-AKT pathway at different levels (Figure 4B). Moreover, among class I of PI3Ks, two out of four members, *Pik3ca* and *Pik3cd,* saw their mRNA expressions respectively 2.5-fold and 5.3-fold-increased in confined K^+^p53^M;T^ spheroids compared to free-growing spheroids (Figure 4C). Only K^+^p53^M;T^ spheroids, harbored a significant 2.2-fold increase in AKT phosphorylation, reflecting the PI3K/AKT pathway activation under confinement (Figure 4D). PI3K/AKT is known to control cell survival (*37, 38*). We thus assessed the level of a cell death marker in the spheroids. K^+^ spheroids displayed a cleaved caspase-3 (cC3) positive core, while K^+^p53^M;T^ showed a more diffuse CC3 distribution (Figure 4E). Confinement of K^+^p53^M;T^ spheroids leads to an early gene response characterized by early plasma-membrane signalling associated with functional activation of PI3K/AKT survival pathway.

**Figure 4.**
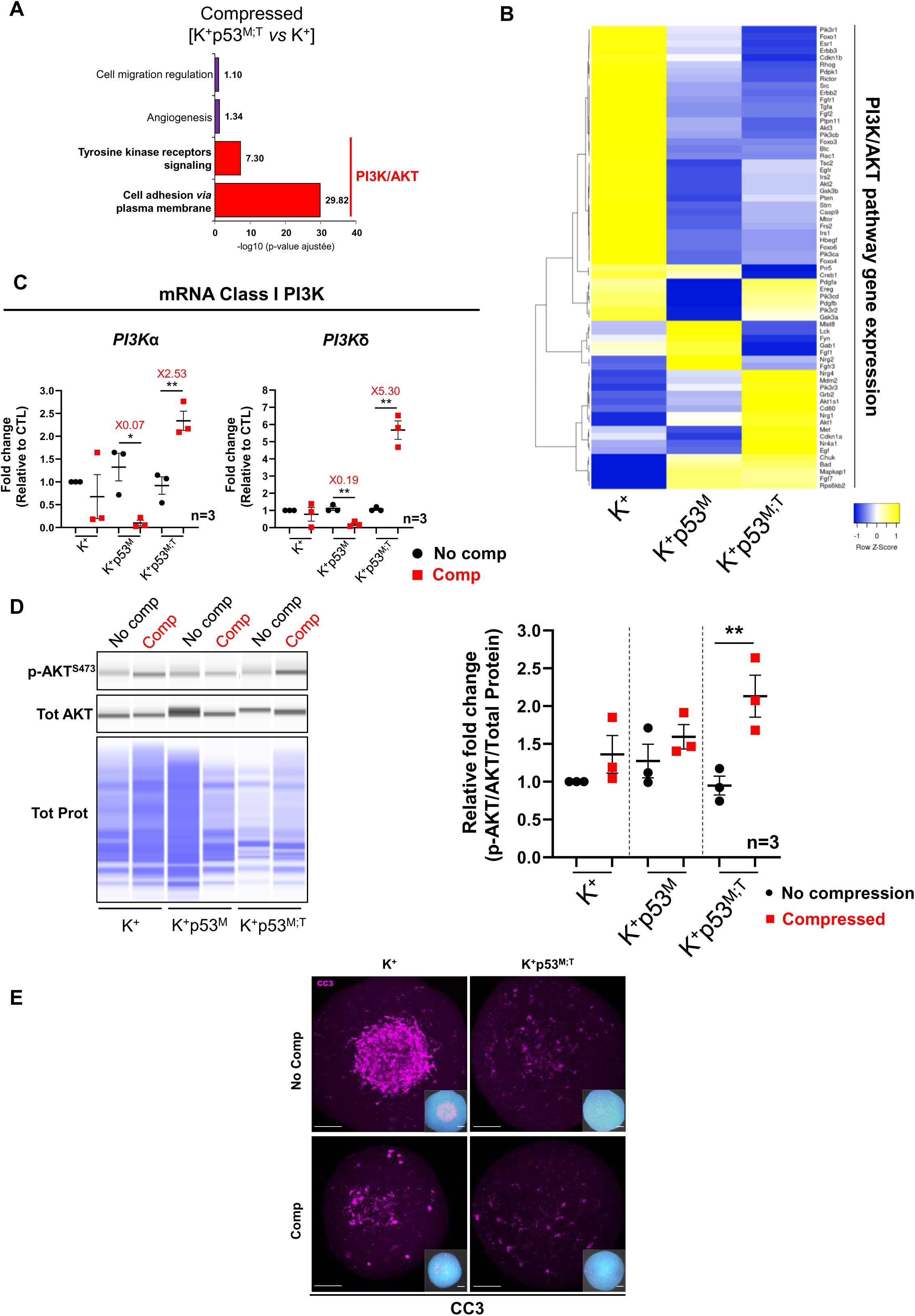
Confinement activates differently PI3K-AKT pathway in mutated-truncated p53 cells. **A.** Gene Ontology analysis shows the enrichment of cellular mechanisms (Tyrosine kinase receptors signaling and Cell adhesion *via* plasma membrane) involving PI3K/AKT pathway in compressed KRAS^G12D^+p53^R172H;R210*^ cells *vs* not compressed KRAS^G12D^+p53^R172H;R210*^ cells. **B.** Heatmap shows the 66 genes of PI3K/AKT signature expression varying depending on the genetic profile under compression in K^+^, K^+^p53^M^ and K^+^p53^M;T^ genotypes. **C.** mRNA expression of PI3K class I members *Pik3ca* and *Pik3cd* in compressed (Comp, red squares) *vs* not compressed (No comp, black dots) in K^+^, K^+^p53^M^ and K^+^p53^M;T^ genotypes. Results are presented as mean, +/- SEM, n=3 x24 spheroids per condition. *p-value<0.05; **p-value<0.01. **D.** AKT pathway activation was analyzed using semi-quantitative measurement of AKT phosphorylation (p-AKT^S473^) related to total AKT (Tot AKT) and total protein (Tot Prot) using simple WB in K^+^; K^+^, p53^M^ and K^+^, p53^M;T^ spheroids (day 6). No comp: No compression; Comp: Compressed. Results are presented as mean, +/- SEM, n=3 x24 spheroids per condition. **p-value<0.01. **E.** Representative maximum intensity projections (MIPs) of confocal optical sections (148–175 planes, depending on spheroid size) showing cleaved caspase-3 (CC3, magenta) and nuclei (DAPI, cyan) in K^+^; K^+^, p53^M^ and K^+^, p53^M;T^ spheroids (day 6), either not compressed (No Comp) or compressed (Comp). Insets display MIPs of cC3 (magenta) together with DAPI-stained nuclei (cyan), allowing visualization of the overall spheroid size and providing spatial context for the localization of CC3 cells within the whole structure. Images were obtained with a Zeiss LSM 780 confocal microscope using a 25× multi-immersion objective (oil/gly ring, NA = 0.8) with laser line 561 nm (CC3) and 405 nm (DAPI). RapidClear 1.49 was used for optical clearing. Scale bar = 100 µm.

### 5. In vivo tumor growth and tumor mechanics interaction differ between the three tested genotypes

We next aimed to test the physio-pathological relevance of our findings in preclinical models of PDAC such as subcutaneous allografts (Figure 5A). *In vivo*, the basal growth rate of K^+^, K^+^p53^M^, K^+^p53^M;T^ tumors was similar (Figure 5B). Fully developed tumors had similar volumes as measured with B-mode echography (Figure 5C). We observed a tendency to an increase of the shear-wave elastography (SWE) measure of the K^+^p53^M;T^ tumor (Figure 5D,E) associated with increased collagen content (Figure 5F, left). The proliferation index as assessed by Ki67 staining and cleaved caspase-3 apoptosis marker remained similar in all tumor genotypes (Figure 5F, center, right), with reduced heterogeneity of the values in K^+^p53^M;T^ tumors. To characterize the mechanical stresses found at tissue level in fully-grown tumors, we adapted a method based on tissue relaxation (Figure 5A) (*39*). The speed and intensity of tissue relaxation after a 4 mm-punch in the center of the tumor was measured through the analysis of the pixel displacement fields at the punch border in a 1400 sec time frame in B-mode echography images using Davis© Software and post-processing routines implemented in python (Figure 5A). We observed that K^+^ tumors accumulated mechanical strains and stresses, as the relative displacement of the punch border was high just after punch compared to K^+^p53^M^ and K^+^p53^M;T^ tumors. In the latter cases, the punch border remained stable in time, as usually measured in healthy tissues (Figure 5G). As this absence of mechanical response was associated with a tendency to an increase of tumor stiffness, as quantified in the pre-punch estimation of an elastic coefficient using shear wave elastography (SWE) (Figure 5D,E), these data highlight that the bidirectional interaction between tumor growth and tumor mechanics could differ between the three tested genotypes.

**Figure 5.**
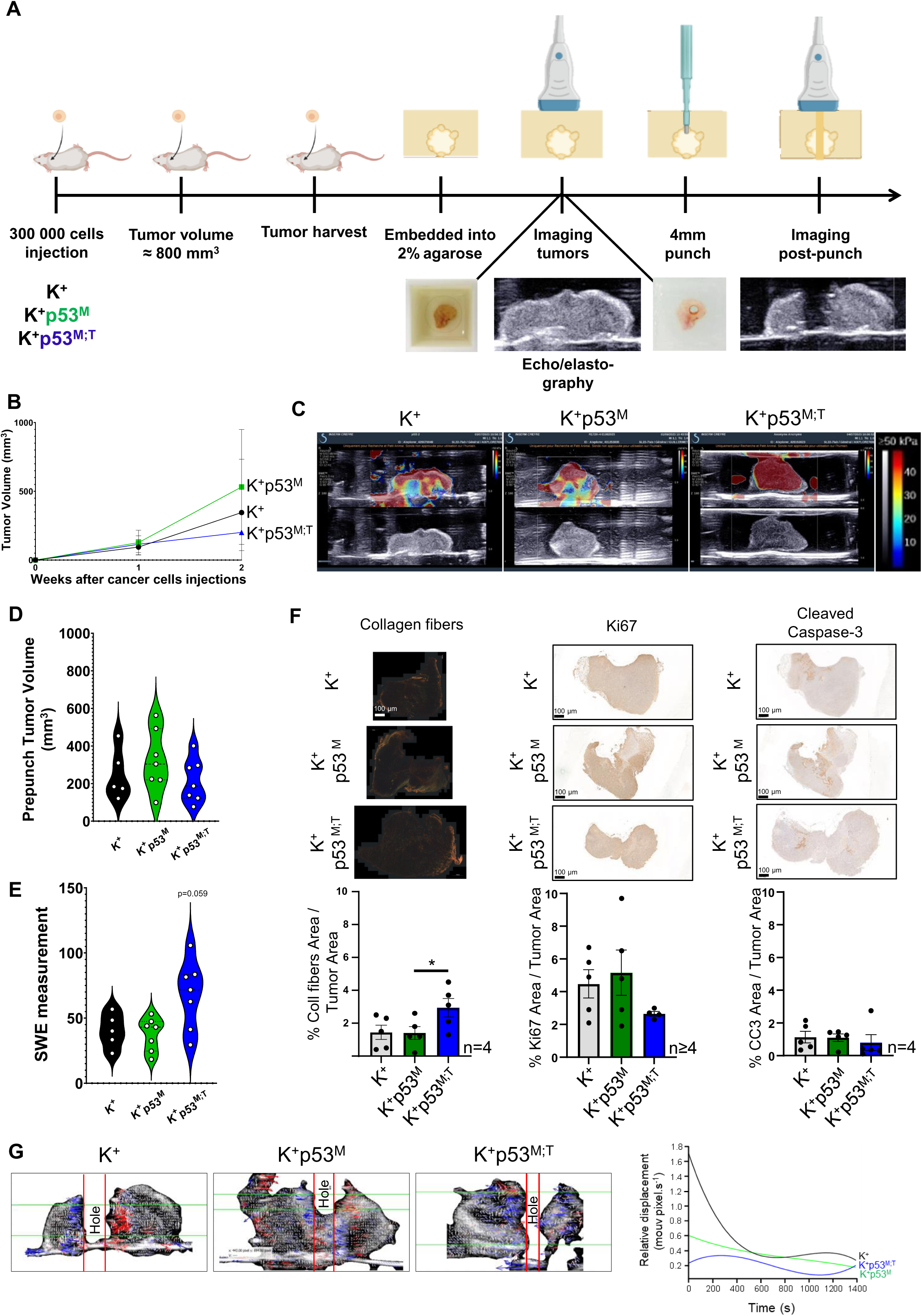
Quantification of mechanical stress at tissular level in fully grown tumors with indicated genotype. **A**. Injection of 300 000 cells in interscapular zone to grow subcutaneous tumors, which were harvested to an equivalent volume. The principle of “hole” or “punch/relaxation” method used in structural mechanics was applied to agarose-coated excised tumors. Tumors were embedded using 2% low gelling temperature agarose as described in (*39*), to stabilise and calibrate the imaging allowing post-processing analysis. In Nia *et al.*(*39*), tumor was then cut to follow stress relaxation using the planar method. Our method based on punch/relaxation method allows the measurement of mechanical stress at tissue level in fully developed tumors. After polymerization of the agarose around the tumor (15 min at 4°C), a hole was made in the tumor (“punch”) using a 4 mm diameter punch. A series of elastography images (Aixplorer) was taken each 30 sec during 1400 sec of tumor relaxation after “punch”. Post-processing of images based on pixel vectorization give access to displacement, strain and stress fields in the tumor during the relaxation of biopsy “hole” to get correlations with cell/tumor genotypes. Post-processing and analysis of the images using COMSOL software v.6.2.0.339 allows us to create a tumor deformation map and infer the associated mechanical stresses to genotype of cells/tumors. **B**. Evolution of tumor volumes (mm^3^) in KRAS^G12D^ (K^+^) (black curve), KRAS^G12D^+p53^R172H^ (K^+^p53^M^) (green curve) and KRAS^G12D^+p53^R172H;R210*^ (K^+^p53^M;T^) (blue curve) tumors in 2 weeks after cancer cells injection. n≥4 tumors per genotype. **C**. Representative images of shear wave elastography in K^+^, K^+^p53^M^ and K^+^p53^M;T^ tumors. Color scale of SWE measurements in kPa. **D**. Tumor volumes estimated with B-mode images taken before punch. Results are presented tumor volumes. n≥4 tumors per genotype. No statistical differences. **E**. SWE measurements taken before punch. Results are presented as mean values measured in the tumoral area. n≥4 tumors per genotype. **F**. Picro Sirius red collagen fibers staining, Ki67 proliferation index and cleaved caspase-3 apoptosis marker pictures and quantifications in KRAS^G12D^ (K^+^), KRAS^G12D^+p53^R172H^ (K^+^p53^M^) and KRAS^G12D^+p53^R172H;R210*^ (K^+^p53^M;T^) tumors. Scale bars represent 100 µm. Results are presented as % of staining area/total tumor area. n≥4 tumors per genotype. *p-value<0.05. **G**. Visualization of tumor relaxation-displacement vectors (up panels) and mean quantification (down panel) of the relative relaxation-displacements of KRAS^G12D^ (K^+^) (black curve), KRAS^G12D^+p53^R172H^ (K^+^p53^M^) (green curve) and KRAS^G12D^+p53^R172H;R210*^ (K^+^p53^M;T^) (blue curve) tumors following the punch/relaxation method.

### 6. In vivo tumor compression shows the importance of mutated-truncated p53 to promote tumor growth

To specifically test the sole effect of compression in a controlled manner *in vivo*, we designed a novel and unique method to compress tumors and mimic a spatial confinement at tissue scale at tumor initiation to test our model (Figure 1A). We developed a minimally invasive compression device using 3D printing. The device imposed unidirectional resulting forces *in vivo*, conferring a localized confinement to the grafted subcutaneous tumor. The location of the subcutaneous injection in the inter-scapular localization of the mouse allowed support of the growing tumor on the rigid spine (Figure 6A; Figure 5; for details of this unique method and device, also see M&M §*In vivo* compression device and associated protocol).

**Figure 6.**
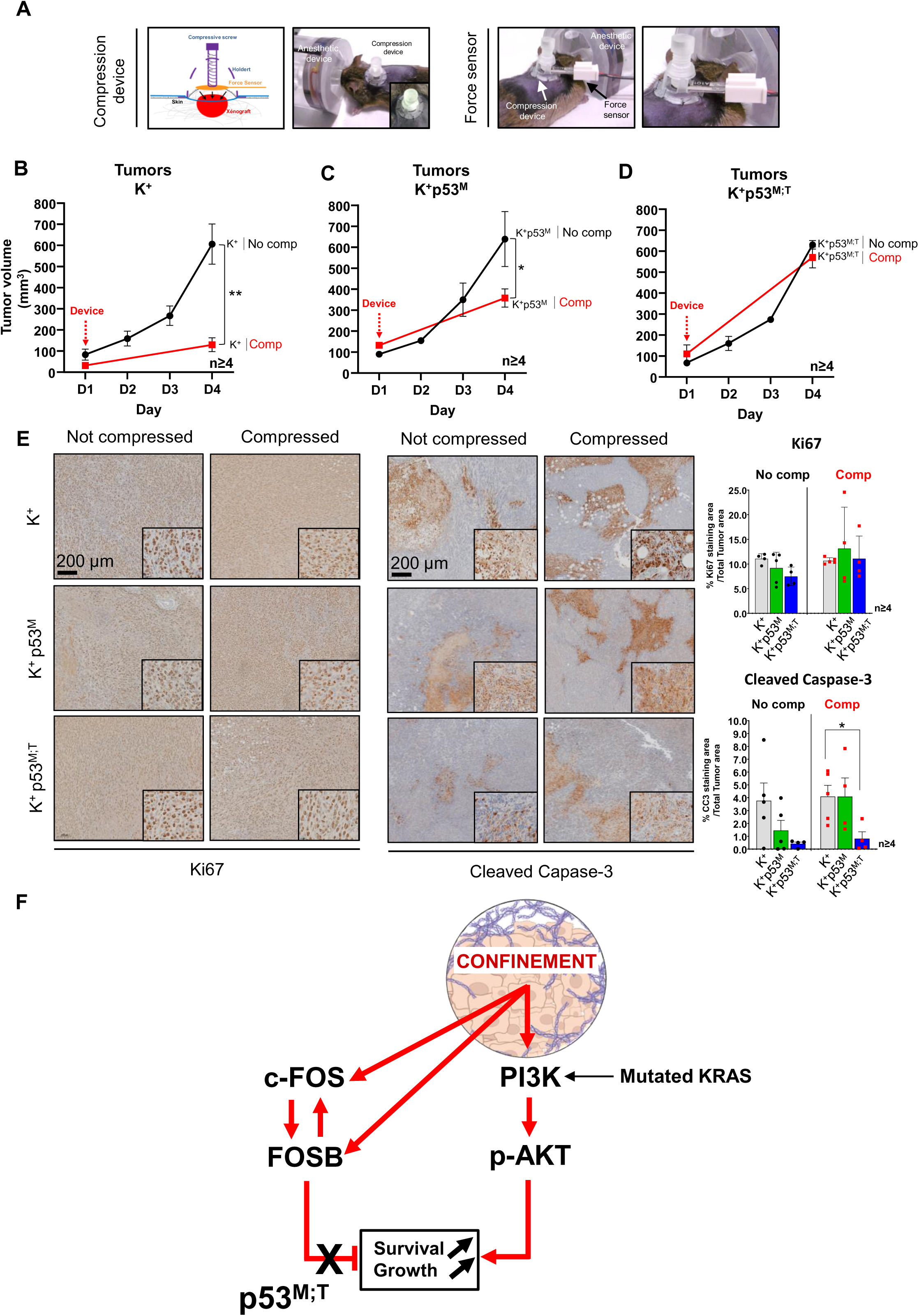
*In vivo* tumor compression shows the importance of p53 double mutant to promote tumor growth, by preventing intratumoral cell death. A. The 3D printed minimally invasive compression device allowing application of *in vivo* allo-xenograft unidirectional force conferring compressive stress to cells (detailed in Supplementary Figure 3A,B) was affixed to the interscapular position of the mice and a force sensor was positioned to calibrate the applied pressure to graft. **B.** K^+^ (left panel), **C.** K^+^p53^M^ (center panel) and **D.** K^+^p53^M;T^ genotype allografts were confined inducing compressive stress (Comp, red curves) compared to not compressed (No Comp, black curves). The tumor volumes were analyzed for 4 days with Day 0 corresponding to compressive device affixation. Results were represented as mean, +/- SEM, n≥4 tumors per condition. *p-value<0.05; **p-value<0.01. **E.** Cleaved caspase 3 and Ki67 staining in 4 days compressed tumors compared to not compressed tumors in K^+^, K^+^p53^M^ and K^+^p53^M;T^ genotypes. The semi-quantitative analysis was performed calculating in % of the cleaved caspase 3 or Ki67 immunostaining area related to the total area of the tumor section. Scale bar corresponds to 200µm. n=4. *p-value<0.05. **F.** Graphical representation of PI3K/AKT pathway activation and FOS (c-FOS and FOSB) transcription factors role as a connected network in p53 mutated truncated condition (p53^M;T^) under confinement in 3D spheroid growth.

The injected isogenic cell lines developed tumors in an interval of 12 to 17 days: K^+^ tumors were detected first (12.7 days ± 1.3 days), followed by K^+^p53^M^ (13.1 days ± 0.8 days) and K^+^p53^M;T^ (17.3 days ± 1.3 days) (Supp Figure 4). Once the tumor reached around 100 mm^3^ diameter, considered as an early stage in tumor growth, the compression mechanism was placed around the tumor, and the tumor was compressed for 4 days. The compression mechanism consisted of a 5 mm diameter holder with a screw entry point on the top and a 10 mm diameter part placed to the skin, surrounding the tumor allograft (Figure 6A; Supp Figure 3). The initial compressive force applied to the tumors was low and calibrated to 0.4 N using a force sensor, which corresponds to 5 kPa of compressive stress (Figure 6A). After 4 days of compression, the growth of KRAS mutated (K^+^) and KRAS and p53 mutated (K^+^p53^M^) tumors was decreased by 79 and 44% respectively, compared to free-growing tumors (Figure 6B-C). However, the volume of tumors harboring p53 mutated-truncated form (K^+^p53^M;T^) remained similar with or without compression (Figure 6D).

Those data clearly indicate a different behavior of tumors, which depends on the genotype of tumor cells. Free-growing mutated-truncated p53 tumors are more rigid, while they continue to grow under confinement. K^+^p53^M;T^ tumors overcome growth inhibition induced by compression, conferring a growing advantage.

### 7. Mutated-truncated p53 tumor growth under compressive stress was associated with cell death

Finally, we aimed to test whether the selective adaptive response to mechanical environment identified in spheroid would be conserved *in vivo* and we used *Fos* gene expression as a proxy of selective adaptive response in K^+^p53^M;T^ tumors. Compressed K^+^p53^M;T^ tumors (Comp) harbored a significant increase in *c-Fos* (3.6-fold) and *FosB* (4.6-fold) mRNA expressions (Supp Figure 5). We previously showed an increase of proliferative and survival PI3K/AKT signal in K^+^p53^M;T^ background. *In vivo*, no difference was found in Ki67 staining across all conditions, indicative of unchanged proliferative index (Figure 6E). We next assessed the level of cleaved caspase 3 cell death marker in the free or compressed tumors. K^+^p53^M;T^ tumors harbored a tendency towards a decrease in cleaved caspase 3 level, indicative of a decreased apoptotic cell death, in uncompressed (No Comp) tumors compared to the other genotypes. This decrease was found significant in compressed (Comp) condition, while cleaved caspase 3 expression was not significantly modified in K^+^ and K^+^p53^M^ tumors with or without compression (Figure 6E). We observed the same cell process as measured in confined spheroids (Figure 4E).

Finally, we evaluated known molecular actors of mechanical stimuli. It was previously shown that CAFs could be responsible for tumor cell compression leading to an inhibition of 3D growth; the inhibition of the transcriptional co-activator YAP was instrumental to this process (*40*). Indeed, tumor compression with the unidirectional compression device was associated with significantly increased the *Yap1* gene expression and of some of its targets (*Ccn1-Cyr61*, *Ap1-c-Jun*), in K^+^p53^M;T^ background only. Other members of YAP pathway (*Ccn2-Ctgf* and *Tead1*) harbored a tendency towards an increased expression in cells with K^+^p53^M^ and K^+^p53^M;T^ genotypes compared to K^+^ cells under compression (Supp Figure 6A).

All these data point to the fact that FOS factors reduced 3D growth under confinement (Figure 6F). However, in K^+^p53^M;T^ tumors, while the expression of FOS factors was significantly increased by confinement, the FOS factor-induced inhibition of 3D growth in this mechanical context was abrogated. Further, PI3K/AKT pathway activation amplified cell survival and growth signaling pathways. Confined K^+^p53^M;T^ tumors are in high oncogenic activation state (high YAP activation) (Supp Figure 6A) and acquire specificities that overcome the anti-growth effect of confinement. These data support the idea that truncated p53 mutants, found in a significant proportion of human tumors, may contribute to cancer progression, particularly in mechanically stressed tumors.

## Discussion

### 1. p53 status and compressive stress in solid tumors

Growth-induced pressure is thought to represent an important determinant of solid cancer progression, particularly in PDAC, a tumor type characterized by an extremely dense stroma (desmoplasia) and high intratumoral pressure (*41–43*). In this study, we demonstrated that p53 genetic background significantly modifies the response of PDAC cells and tumors to confinement. This finding shows that the genetic environment, particularly alterations in *TP53*, is decisive in the response of cancer cells to mechanical loadings. Whereas wild-type p53 cells display reduced viability under confinement, cells harboring a mutated-truncated p53 protein exhibit growth maintenance under confinement, highlighting a critical role of p53 status in mechano-adaptation. These observations are consistent with earlier reports that compressive stress modifies cell proliferation and death, collapses blood vessels, and drives necrotic core formation in pancreatic and breast cancers (*44–46*). Compressive stress was found to promote mitochondria-dependent cell death (*47*). Blood vessel collapse induces hypoxia mechanism and induces HIF1α expression and accumulation in tumor cells (*48*). However, here 3D tumor compression did not significantly affect *Hif1α* gene expression in the three different genotypes with or without compression (K^+^, K^+^p53^M^, K^+^p53^M;T^) (Supp Figure 6B). This result could be explained by the fact that pancreatic tumors are poorly vascularized (*49*) and hypoxia could not be further modulated by compression.

In addition to pancreatic and breast cancers, compressive stress is high in colon cancer and associated with genetic alterations such as KRAS and p53 mutations (*50–54*). This solid cancer develops through a multistep process initiated by specific mutations in proto-oncogenes and tumor suppressor genes (such as *KRAS* and *TP53*) (*55*). In addition, in this cancer, mechanical stimuli can initiate tumorigenesis by modulating the WNT pathway and promote metastasis by driving phenotypic shifts and mechanical adaptations that enable tumor cells to survive intravasation, circulation, and extravasation (*52*).

Moreover, alterations in mitochondrial apoptosis regulation emerge as a key converging point between AP-1 protein activity and the gain-of-function properties of truncated p53 mutants (*56, 57*). c-FOS and FOSB influence mitochondrial apoptotic sensitivity through AP-1–dependent induction of anti-apoptotic BCL-2, a mechanism shown to preserve outer-membrane integrity and limit cytochrome-c release (*58, 59*). In parallel, exon-6 truncating p53 mutants generate a diverse function isoform of p53 described by Shirole *et al*. (*27*), which not only lose canonical pro-apoptotic transcriptional activity but also acquire mitochondrial functions that decrease cell death. Notably, the exon-6 p53 truncated forms modulate cyclophilin D (CypD), a key regulator of the mitochondrial permeability transition pore (mPTP), reducing its ability to promote mitochondrial depolarization and permeability, and thereby further restricting apoptosis (*27*). These effects are reinforced by the stabilization of anti-apoptotic BCL-XL and MCL-1 at the outer membrane (*60, 61*). Altogether, the anti-apoptotic effect of exon-6 truncated p53 isoforms and AP-1 members associated with pro-proliferative effect of PI3K-AKT signaling amplifies the cell pro-survival program that consolidate cell resistance to apoptosis and promote pro-proliferative behavior (*27, 62, 63*).

Our findings extend our knowledge by demonstrating that genetic alterations in *TP53* gene can fundamentally reshape pathways involved in mechanosensing, change tumor cell fate to adapt and resist compression stress, thereby influencing solid cancer progression.

### 2. p53 genetic alterations and mechanotransduction

Genetic alterations intersect with mechanotransduction pathways to shape tumor progression. In colorectal and pancreatic cancers, mutations in oncogenes such as *KRAS* and *PIK3CA*, or in tumor suppressors including *TP53* and *APC*, not only reprogram classical signaling cascades but also enhance cellular responsiveness to mechanical stress within the tumor microenvironment (*55, 64–66*). Mechanical cues such as compressive stress, shear stress and tissue stiffening have been shown to activate pro-survival pathways like transcriptional regulators such as YAP/TAZ, thereby promoting proliferation, stemness, and resistance to apoptosis (*44, 67*). Notably, in colorectal cancer, the protein VASN amplifies tumor progression by engaging both YAP/TAZ and PI3K/AKT pathways; VASN interacts with YAP to suppress its inhibitory phosphorylation and concurrently activates the PI3K/AKT axis, enhancing proliferation, invasion, and EMT (*68*). In our previous study, we did not observe any modification of YAP expression in p53 untruncated pancreatic and breast cancer cells after 24h of 2D compressive stress (*10*). However, YAP/TAZ Hippo and non-Hippo pathways are known as key regulators of mechanotransduction under mechanical stress (*69*). Here, we found an overexpression of *Yap1* gene and overexpression of downstream targets of YAP (*Ccn1-Cyr61*, *Ap1-c-Jun*) and this was associated with a strong activation of PI3K/AKT pathway depending on the p53 mutational background of tumors. This finding is in line with previous studies involving PI3K/AKT pathway with mechanical compressive stress response in pancreatic and breast cancer cells (*10, 38*). Beyond survival under such mechanical stimuli, this study revealed that mutated–truncated p53 cells exhibited a high activation state and unique phenotypic specificities under mechanical stress. Notably, it strongly activated the PI3K–AKT pathway in truncated p53 cells, linking stress with intra-tumoral pro-survival signaling. Previous studies have shown that compressive stimuli could stimulate migration and activate oncogenic cascades such as AKT/CREB1 in pancreatic cancer (*42*).

Importantly, the mutational background determines how cells integrate these mechanical signals: for instance, truncated or mutated p53 can impair the effect of the early gene transcriptional responses by AP1 factors family (c-FOS and FOSB) while enabling adaptive oncogenic pathways such as PI3K/AKT and YAP. This interplay may explain why *in vivo* tumors harboring truncated p53 displayed reduced cell death and greater growth maintenance under compression, ultimately conferring a growth advantage in mechanically hostile microenvironments. These observations highlight a synergistic link between genetic lesions and mechano-transduction, positioning mechano-genetic interactions as critical determinants of tumor aggressiveness and potential therapeutic targets. To go further, future studies should therefore explore the mechano-genetic interplay in more complex carcinogenesis models, integrating single-cell analyses to dissect heterogeneity in response to compressive stimuli.

This study proposes that cells carrying truncated p53 showed a remarkable ability to withstand compressive stresses compared to their wild-type counterparts. At long term, our findings raise opportunities for solid cancer therapeutic targeting. As targeting mechano-sensitive pathways such as PI3K–AKT may represent a promising therapeutic avenue in patient’s care, p53 mutational status could serve as a biomarker for predicting tumor adaptation to mechanical stress and efficiency of mechanotherapies to be developed in the future (*70*).

## Materials and Methods

### Plasmids, sub-cloning and mutagenesis

*p53^R172H^* and *p53^R172H;R210*^* cDNAs were amplified by polymerase chain reaction (PCR) using CloneAmp™ HiFi PCR Premix (Takara Bio Inc, Shiga, Japan) according to the manufacturer’s protocol, primers listed in Supporting Table 2 and sub-cloned in pSBtet-GFP-Neomycin (pSBtet-GN; Addgene plasmid # 60501; a gift from R. Marschalek (*71*) using engineered SfiI cloning sites. *p53^R172H^* mutation and *p53^R172H;R210*^* mutation and stop codon were inserted using the QuikChange II XL mutagenesis kit (Agilent, Santa Clara, CA, USA) and the primers listed in Supporting Table 2.

Amplicons were then digested 30 min to 1 h at 50°C with SfiI (NEB Ipswich, MA, USA). The ligations were performed 30 min at room temperature using DNA Ligation Kit, Mighty Mix (Takara Bio Inc, Shiga, Japan). pCMV(CAT)T7-SB100 plasmid coding for Sleeping Beauty Tranposase (Addgene plasmid # 34879) was a gift from Z. Izsvak (*72*). All the plasmid constructions were validated by sequencing using primers listed in Supporting Table 2.

### Cell culture, transfections and stable cell lines generation

Pancreatic cancer cells (KRAS^G12D^ mouse PDAC derived cells (*73*) and induced Mouse Embryonic Fibroblasts (iMEF)(*74*) were cultured in Dulbecco’s Modified Eagle Medium (DMEM) (GIBCO; #61965026) supplemented with 10% fetal bovine serum (Eurobio Scientific; #CVFSVF00-01), Penicillin-Streptomycin 50 U/mL and 50 µg/mL respectively (Sigma-Aldrich; #P0781), L-Glutamine 2 mM (Sigma-Aldrich; #G7513) and Plasmocin 25 µg/mL (InvivoGen; #ant-mpp) and maintained at 37°C in humidified atmosphere with 5% CO2. For transient transfections, cells were grown in 60 mm dishes and transfected with 3 µg plasmid, using jetPRIME® (Polyplus-Transfection, Illkirch-Graffenstaden, France) for 24 h. DNA was transfected 24 h after plating the cells on 60 mm dishes, and the medium was changed every day. For stable transfections, pancreatic cells were grown in 35 mm dishes. Cells in each well were transfected with 1.9 µg plasmid (pSBtet-GN, pSBtet-GN-p53^R172H^ or pSBtet-GN-p53^R172H;R210*^) and 100 ng of pCMV(CAT)T7-SB100 vector, using jetPRIME® (Polyplus-Transfection, Illkirch-Graffenstaden, France). Twenty-four hours after transfection, cells were treated with 1 mg/mL G418 for 7 days. GFP-positive cells were sorted using LSR Fortessa™ X-20 Cell Analyzer (BD, Franklin Lakes, USA) and distributed one cell per well for clonal selection. After growth for 5 days in DMEM, 10% FBS, 2 mM L-Glutamine, Penicillin-Streptomycin (50 U/mL and 50 µg/mL), each clone was treated with 1 mg/mL G418 for 2 weeks and the stability of the transfected vectors was monitored by detecting GFP fluorescence. Cells were treated with 0.5 µg/mL doxycycline for 5 days to induce p53^R172H^ and p53^R172H;R210*^. KRAS^G12D^ mutated cells were annotated in the text: K^+^; KRAS^G12D^p53^R172H^ annotated: K^+^p53^M^ and KRAS^G12D^p53^R172H;R210*^ annotated: K^+^p53^M;T^ (Figure 1B, Supp Figure 1).

### Normalized cell number assay

Cells were rinsed with PBS (Eurobio Scientific, #CS1PBS 0101) and fixed for 15 min using PBS containing 10% methanol and 10% acetic acid. Cells were stained for 15 min using crystal violet (Sigma Aldrich; #HT90132). Images were taken using Chemidoc™ Imaging System (BioRad) and quantified using ImageJ software.

### Spheroids generation and agarose confinement

1x10^3^ KRAS^G12D^, KRAS^G12D^p53^R172H^ and KRAS^G12D^p53^R172H;R210*^ PDAC cells were mixed in 10 µL droplets affixed on 100 mm petri dish cover using DMEM (GIBCO; #61965026) supplemented with indicated products in §*Cell culture, transfections and stable cell lines generation* and 0.5 µg/mL doxycycline for 2 days. The petri dish cover was flipped over in order to allow aggregation of cells in a meniscus formed by the hanging drop. The 100 mm petri dish was filled with 20 mL Phosphate Buffered Saline (PBS) (Sigma-Aldrich; #D8537). After 48h, spheroids were formed and harvested. Half of them were embedded in 1% low gelling temperature agarose (Sigma-Aldrich; #A0701) in Phosphate Buffered Saline (PBS) containing 0.5 mM MgCl_2_ and 0.9 mM CaCl_2_ respectively (Sigma-Aldrich; #D8662) to confine it and induce a growth-induced stress on cells and the other half also called group of free spheroid or no comp was deposited on 1% low gelling temperature agarose without compressive stress. Knowing the physicochemical characteristics of 1% low gelling temperature agarose, the growth-induced pressure was estimated using the following formula: (-5.772*10^-5^*100*(R(tx)-R(t0)/R(t0))^4^ + (0.01048*100*(R(tx)-R(t0))/R(t0))^3^ + (-0.9048*100*(R(tx)-R(t0)) / R(t0))^2^ + (56.25*100*(R(tx)-R(t0)) / R(t0)) + 0.03247 with R(tx)= spheroid radius at t (day 2, day 4 or day 6) and R(t0)= spheroid radius at day 0 (*75, 76*). This formula was validated for early growth (when spheroid radius increases by less than 50%). Day 0 was considered at 1% low gelling temperature agarose spheroid embedded day (Supp Figure 1D). All confined and not confined spheroids were cultured in DMEM, 10% FBS, 2 mM L-Glutamine, Penicillin-Streptomycin (50 U/mL and 50 µg/mL) with 0.5 µg/mL doxycycline. 1% low gelling temperature agarose is permissive to medium substances and oxygen (*77, 78*). No necrotic cores were observed analyzing spheroids in brightfield. All these spheroids were used to analyze spheroid growth (radius measurement), western blot, RNA sequencing, RT q-PCR.

### siRNA transfections in spheroids

Spheroids were performed as described in § *Spheroids generation and agarose confinement*. During spheroids formation, cell droplets were transtected with 10 nM siRNA (siRNA scramble, siRNA against *c-Fos*, siRNA against *Fosb* or siRNA against *c-Fos* and *Fosb* described in Supporting Table 2) using jetPRIME® (Polyplus-Transfection, Illkirch-Graffenstaden, France) for 48 h. After spheroid formation, they were maintained in 10 nM siRNA medium or in 1% agarose + siRNA medium for 6 days of spheroid growth. To maintain 10 nM siRNA concentration along the experiment, medium was replaced by fresh medium + siRNA each 48 h.

### Protein extraction from cells and spheroids

For one loaded sample, 12 spheroids were harvested in 15 mL tube. For embedded spheroids, 1% agarose was dissolved using solubilisation buffer (QiAquick®, Qiagen, Hilden, Germany) for 10 min at 50°C. Spheroids were spined down in a 15 mL tube and supernatant was removed. Spheroids were lysed and resuspended in a lysis buffer containing 150 mM NaCl, 50 mM Tris, 1 mM EDTA, 1% Triton, 2 mM dithiothreitol (DTT), 2 mM Sodium fluoride, 4 mM Sodium Orthovanadate and supplemented with protease inhibitors (Complete protease inhibitors, Roche). After a 20 min incubation on ice, a 10 min centrifugation was performed at 12,000g and 4°C and the supernatant was collected. Protein concentration was measured using Bicinchoninic assay (BC Assay Protein Quantification Kit, #3082, Interchim). For 2D cells, cells were rinsed with PBS and detached by scrapping in PBS. Cells were harvested through a 5min centrifugation at 3,000g and 4°C and lysis buffer was added in cells as above protocol.

### Western Blot and simple Western Blot

20 µg of proteins were separated on a 10% polyacrylamide gel and transferred on nitrocellulose membranes (Amersham™ Protran®; #106000004) using Trans-Blot® Turbo™ (BioRad). Blocking was performed through a 1h incubation in 5% low fat milk in TBST. Membranes were incubated overnight at 4°C with primary antibodies in TBS-Tween 0.1%, 5% Bovine Serum Albumin as indicated in Supporting Table 2. Then, membranes were rinsed three times with Tris-buffered saline 0,1% Tween (TBS-Tween 0.1%), incubated for 1h with secondary antibodies coupled to a horseradish peroxidase, in 1% low fat milk described in Supporting Table 3. Membranes were next washed three times (1 min, 5 min and 10 min) with TBS-Tween 0.1%. Proteins were detected through chemiluminescence (Clarity™ Western ECL Substrate; BioRad; #1705061) using Chemidoc™ Imaging System (BioRad) and quantified using ImageJ software. ≥3 independent replicates were performed for each protein analyzed.

For simple western blot, 2 µg of proteins were separated on a 10% polyacrylamide gel. Samples were prepared according to the manufacturer recommendations: 12-230 kDa Separation Module Catalog #SM-W001 (Protein Simple-Bio-Techne, Minneapolis, MN, USA). Briefly, standard pack reagents were prepared (400 mM DTT, 20µL of 10X Sample buffer and 20 µL of 400 mM DTT in 5X Fluor master mix, and 20 µL of water in ladder). Samples were prepared using 5X Fluor master mix and gently mixed by up and down pipetting. They were denatured 5 min at 95°C, vortex and spin down. Reagent, samples and primary/secondary antibodies (listed in Supporting Tables 2-3) were loaded in Jess Automated Western Blot System® (Protein Simple-Bio-Techne, Minneapolis, MN, USA) and results were analyzed using Compass software (Compass Software GmbH, Dortmund, Germany). Signal linearity in JESS was in the range where the detected signal was directly proportional to the protein amount, ensuring accurate and reliable quantification.

### 2D Immunostaining and microscopy

Cells were cultured on glass coverslips. After compression (described in (*10*)), cells were fixed with 4% PFA in PBS for 10min and then permeabilized with 0.1% Triton in PBS for 5min. Cells were blocked in blocking solution (1% BSA in PBS) for 30min. Samples were incubated with p53/GFP primary antibodies (Supporting Table 2) diluted into blocking solution overnight at 4°C. Cells were washed with PBS and then incubated with the Alexa Fluor® 488 secondary or Alexa Fluor® 594 antibodies diluted into blocking solution for 1h (Supporting Table 3). Samples were washed with PBS and incubated with DAPI (Sigma; #D9542 0.1µg/ml) as nuclear counterstain, for 3 min. Coverslips were mounted in Fluoromount-G (Invitrogen; # 00-4958-02). Images were acquired with a Plan Aprochromat 63x ON 1.4 oil immersion objective using a Zeiss LSM780 confocal Microscope using Airyscan with post-processing orthogonal projection.

### 3D Immunostaining and imaging

An optimized immunofluorescence workflow on 3D spheroids was developed. Spheroids (n =12 per conditions) were washed twice in PBS. Fixation was performed in freshly prepared 4% paraformaldehyde (PFA) for 1 h 30 min at room temperature under gentle agitation. Permeabilization was achieved with 0.5% Triton X-100 in PBS for 30 min, followed by blocking for at least 2h in 3% BSA containing 0.2% Triton X-100. All steps were carried out at room temperature on a tilted metallic support to facilitate uniform exposure. Primary antibody incubation was optimized for each target (antibody titration, incubation time, and buffer composition). Primary antibodies (listed in Supporting Table 3) were diluted in antibody buffer (3% BSA + 0.1% Triton X-100) and incubated for 2 days at 4 °C under gentle agitation on an inclined metallic support to ensure homogeneous antibody penetration. All experimental groups, 2 spheroids for each (not compressed vs compressed; K+ vs K+p53M; T) were processed in parallel with same protocol conditions to ensure comparability. After primary incubation, spheroids were washed three times in 1% BSA + 0.1% Triton X-100 in PBS, with a final 20 min wash in 0.1% Triton PBS under agitation. Residual buffer was carefully aspirated before proceeding to secondary labeling. Secondary labeling was performed in antibody solution (1% BSA + 0.1% Triton X-100 in PBS) containing rabbit AF568 and DAPI (Supporting Tables 3,4), incubated overnight at 4 °C under gentle agitation on a tilted metallic support. Stained spheroids were washed in PBS extensively before clearing and mounting. Clearing efficiency was strongly dependent on spheroid size and intrinsic cell line-specific properties, notably extracellular matrix content. Post-fixation before clearing improved staining preservation. Spheroids were post-fixed with fresh prepared 4% PFA for 30 min, washed twice in PBS, and cleared with RapiClear 1.49 (SUNJin Lab, Taiwan) for at least 24 h. Samples were mounted using dual spacers (0.25 µm) (SUNJin Lab, Taiwan) to avoid deformation, and very importantly with refractive index matched to the objective immersion medium to optimize depth imaging.

We also performed microscopy and workflow optimization. 3D imaging was performed using a Zeiss LSM780 confocal microscope equipped with a 25× long-working-distance multi-immersion objective (NA 0.8) with adjustable correction collar (“multi-immersion ring”). Z-stacks were acquired every 2.8 µm using sequential scanning of each channel. Excitation was achieved with **405 nm (DAPI)** and **561 nm (AF568)** lasers. GaAsP detector gain and emission windows were optimized to prevent cross-talk and saturation. Z-compensation was applied to preserve signal uniformity with depth, according to Nürnberg *et al.*, 2020 (*79*).

### RNA extraction

For RNA extraction, 12 spheroids were harvested for each replicate in each condition in 15 mL tube. For embedded spheroids, 1% agarose was dissolved using solubilisation buffer (QiAquick®, Qiagen, Hilden, Germany) for 10 min at 50°C. Spheroids were spined down in a 15 mL tube and supernatant was removed. RNA was extracted using RNAqueous™-Micro Total RNA Isolation Kit #AM1931 (ThermoFisher Scientific, Waltham, MA, USA) according to the manufacturer recommendations. Briefly, cells were lysate using lysis buffer from Isolation Kit #AM1931 for 5 min at 4°C temperature and lysate was loaded in RNA collection column and centrifuge at 10000g for 1 min at 4°C. Column were rinsed and centrifuge at 10000g for 1 min at 4°C two times using wash buffer from Isolation Kit #AM1931. Elution buffer from Isolation Kit #AM1931 was added in the RNA collection column for 5 min at 4°C. Column were inserted in collection microtube and centrifuge at 10000g for 1 min at 4°C. RNA was collected in the collection tube and RNA concentration was measured using NanoDrop™ 2000 (ThermoFischer scientific, France). Quality of samples was controlled by NanoDrop (NanoDrop 2000 (Thermo Fisher Scientific), Qubit fluorometry (Invitrogen, Carlsbad, CA)) and Fragment Analyzer (Agilent, Les Ulis, France) for measuring concentration and calculation of RNA integrity number (RIN). Samples with RIN<7 were not sequenced.

### Reverse transcription and quantitative PCR

cDNA synthesis was performed with iScript kit (BioRad; #1708891) according to the manufacturer protocol using C1000 Touch Thermal Cycler (BioRad) and the following conditions: annealing: 5 min at 25°C, reverse transcription: 20 min at 46°C, reverse transcriptase inactivation: 1 min at 95°C. cDNAs were diluted to a half in RNAse free water. Gene expression was quantified with the SsoFast EvaGreen supermix kit (BioRad, #1725204) using a thermocycler (StepOne™ #4376374; Software v2.2.2) with the following conditions: 20 sec at 95°C, 40 denaturation cycles: 3 sec at 95°C, annealing and elongation: 30 sec at 60°C. β-actin was used as housekeeping gene. The primers used are described in Supporting Table 1 and each amplicon was validated by sequencing. Gene expression quantification was performed using the Livak method: 2^-(ΔΔC(T))^ (*80*). All RT-qPCR fragments were validated by sequencing using primers listed in Supporting Table 2. Distinct RNA samples ≥4 were analyzed, each amplification was performed in technical duplicate.

### RNA sequencing and bioinformatics analysis workflow

RNA was extracted using previous protocol (§*RNA extraction*). RNA sequencing was performed by the Next Generation Sequencing Service (NGS, Azenta Life Sciences (Burlington, MA, USA)). Quality of samples was controlled by NanoDrop (NanoDrop 2000 (Thermo Fisher Scientific), Qubit fluorometry (Invitrogen, Carlsbad, CA)) and Fragment Analyzer (Agilent, Les Ulis, France) for measuring concentration and calculation of RNA integrity number (RIN). Samples with RIN<7 were not sequenced.

cDNA libraries were sequenced using Illumina NovaSeq, PE 2x150 sequencing platform and configuration with estimated data output of 12 G of raw data per sample.

Bioinformatics analysis were performed by Azenta Life Sciences (Burlington, MA, USA). Read quality control was performed using FastQC software v.0.11.9 (Babraham Institute, Cambridge, UK). Sequence reads were trimmed to remove adapter sequences and nucleotides with poor quality using Trimmomatic v.0.36 (RWTH Aachen University, Germany). The trimmed reads were mapped to the *Mus musculus* GRCm38 reference genome (ENSEMBL) using the STAR aligner v.2.5.2b. The STAR aligner is a splice aligner that detects splice junctions and incorporates them to help align the entire read sequences. BAM files were generated as a result of this step. Unique gene hit counts were calculated by using featureCounts from the Subread package v.1.5.2. The hit counts were summarized and reported using the gene_id feature in the annotation file. Only unique reads that fell within exon regions were counted. After extraction of gene hit counts, the gene hit counts table was used for differential expression analysis. A comparison of gene expression between the customer-defined groups of samples was performed using DESeq2 v.1.40.2. The Wald test was used to generate p-values and log2 fold changes. Genes with an adjusted p-value < 0.05 and absolute log2 fold change > 1 were called as differentially expressed genes for each comparison. The results of the number of significantly differentially expressed genes for all comparisons were provided. Data are stored in an open access repository (accession number available upon request).

A gene ontology analysis was performed on the statistically significant set of genes by implementing the software GeneSCF v.1.1-p2. The Gene Ontology (GO) list was used to cluster the set of genes based on their biological processes and determine their statistical significance. The p-value in GO enrichment analysis represents the probability that the observed overrepresentation of a GO term within a gene set occurs. A low p-value indicates significant enrichment of the GO term. A list of genes clustered based on their GO was generated.

### *In vivo* compression device and associated protocol

Our lab developed a 3D printed minimally invasive compression device compatible with animal experimentation. It allowed the application of *in vivo* allo/xeno-graft unidirectional stress at a given unit calibrated using the force sensor and conferring compressive stress to allo/xenograft tumors. This device is uniquely available and easy-to-produce for compressing subcutaneous xenografts. It is adaptable to diverse subcutaneous allo/xeno-grafts. This device was 3D printed using a biocompatible, transparent and gas permeable PDMS (poly-di-methyl-siloxan) polymer and is composed of a 20 mm length, 5 mm large screw entering into a 5 mm diameter holder for screw entry point on the top and 10 mm diameter part placed on the skin, encompassing the tumor allo/xenograft (Supp Figure 3A,B). Associated to the device, we developed a compression protocol associated: 300,000 mouse pancreatic cancer cells were injected subcutaneously in the interscapular localization of Nude mouse (location allowing support on the spine). Injection of 300,000 cells enables daily monitoring of tumor growth without rapidly exceeding the device holder’s maximum size and the ethically acceptable volume of 1000 mm³. Once the cells were injected, the tumor growth was controlled using tumor volume and the compression device was affixed by 8 stitches on the mouse skin at 100 mm^3^ (± 20 mm^3^). The tumor was compressed for 4 days (Supp Figure 3B). The compression at Day 0 (day of initial compression) was set at a fixed value using a force sensor (FlexiForce® OEM Kit, Teckscan, South Boston, MA, USA) apposed between the subcutaneous tumor and the screw. The force sensor was previously calibrated using a series of known masses to ensure accurate and reliable force measurements. Moreover, mice tumors were visually controlled every day for 4 days and volumes were measured at Day 0 and Day 4 (Supp Figure 3B). In parallel, 300,000 mouse pancreatic cancer cells were injected subcutaneously in the interscapular localization of the mouse, and tumor volume was measured each day during 4 days and compared to compressed allograft tumor. Tumor volume was assessed using this formula V= l²xLx0.52. After compressive device was removed and tumors were harvested for analysis.

### Quantification of mechanical stresses accumulated during tumor development using the “punch/relaxation” method

Inspired by the principle of “hole” or “punch/relaxation” method used in structural mechanics (ASTM E837-20), we designed a protocol for biological objects. To ensure reliable measuring, standardisation of the method was developed as described in Marty *et al.*, in preparation. This method can be applied to PDAC tumors and also to other types of tumors allo/xeno-graft in mouse. 300,000 mouse pancreatic cancer cells harboring the different genotypes K^+^, K^+^p53^M^ and K^+^p53^M;T^ were injected subcutaneously in the interscapular localization of NSG mice. After 21-28 days, tumors were dissected -with average similar volumes in each group (maximal volume was 563 mm^3^) measured with non invasive echography imaging (Aixplorer®, Supersonic, Paris, France). These tumors were embedded using 2% low gelling temperature agarose (Sigma-Aldrich; #A0701) as described in (*39*). After polymerization of the agarose around the tumor (15 min at 4°C), a hole was made in the tumor (“punch”) using a 4 mm diameter punch (#49401, Pfm medical, Merignac, France). Before the “punch”, a mode B ultrasound and an elastography shear wave measurement (Aixplorer®, Supersonic, Paris, France) were performed. A series of mode B ultrasound images was then taken each 30 sec during 1400 sec of tumor relaxation after “punch”. Post-processing of images based on pixel vectorization (Davis© softaware and python scripts) gives access to displacement, strain and stress fields in the tumor during the relaxation of the biopsy “hole” and provided correlations with tumor genotypes. Specific software was developed to help classification and data storage of images; it is available here: https://archive.softwareheritage.org/browse/directory/c7411303bd02da1a8558ca3d49b7eee0e9e81baf/?origin_url=https://github.com/FredPont/remove_subdirs&revision=00f3d00ff74bfef6f1ba8ca68af39c6b109a2900&snapshot=61ac07b36df6deb7a0b345c6c75a5f3eb00f3c51 https://github.com/FredPont/remove_subdirs/releases/tag/v20250430.

### Immunohistochemistry

Immunohistochemistry was performed on 4 µm sections of formalin-fixed paraffin-embedded tissues. Briefly, tissue sections were deparaffinized 3x 5 min in xylene, 3x 5 min in 100% and 1x in 70% ethanol and washed 1x in tap H_2_O and 1x in deionized H_2_O. The antigen unmasking was performed using autoclave heating at 121°C, 250 bar for 12 min (Advantage-Lab, AL02-03) in 100 mM sodium citrate pH 6.0. Sections were blocked for 60 min at room temperature in humid chamber using 2.5% horse serum blocking solution (Vector Lab, MP-7401)). Primary antibodies were diluted in antibody diluent (Dako REAL AG-S202230-2) and incubated at 4°C overnight in humid chamber. Sections were blocked using endogenous peroxidase in PBS + 3% H_2_O_2_ for 10 min at room temperature. After, anti-rabbit Immpress solution (Vector Lab, MP-7401) was used as secondary antibody detection for 60 min at room temperature in humid chamber. Antibody staining was revealed using DAB peroxidase substrate (Vector Lab, SK-4105) respectively 30 sec for Ki67 and 60 sec for CC3 staining. Mayer’s hematoxylin (Merck, 109249) counterstain was performed for 60 sec, following by washes in tap H_2_O. Sections were briefly rinsed in 0.1% HCl and washed using tap H_2_O. Coverslips were placed on slides using Glycergel solution (Dako, C0563). Pictures were taken using Axioscan 7 imager (Carl Zeiss, Germany) and observed using NDP software (San Francisco, USA). Quantifications were performed using QuPath software (*81*), using Pixel classifier tool. The following parameters were used: for Ki67, Resolution: very high, Sigma: 0-3, Threshold: 0.23-0.45, Above: Ki67, Below: Negative; for For CC3: Resolution: very high, Sigma: 0-0.5, Threshold: 0.25-0.5, Above: CC3, Below: Negative. Negative controls were used to validate specific stainings. Primary and secondary antibodies were described in Supporting Tables 3 and 4.

### Picro Sirius red coloration

Picro Sirius red coloration were performed on 4 µm sections of formalin-fixed paraffin-embedded tissues. Briefly, tissue sections were deparaffinized as described above, followed by Picro Sirius red solution (Abcam, ab15068, Cambridge, UK) incubation for 30 min at room temperature. After sections were 2x briefly washed in 0.5% acetic acid, after dehydrated in 95% ethanol for 2 min, 100% ethanol and xylene for 5 min. Coverslips were placed on slides using Eukitt solution (Sigma, 03989). Pictures were taken with Axioscan 7 imager (Carl Zeiss, Germany) using circular polarized light and observed using Zen software (Carl Zeiss, Germany). Quantification were performed using QPath software(*81*), using Pixel classifier.

### Animals

All animal procedures were conducted in compliance with the Ethics Committee pursuant to European legislation translated into French Law as Decret 2013-118 dated February, 1^st^ 2013 (APAFIS#2020101311044955). Grafts were performed in interscapular localization of male Swiss Nude mice (Charles River Laboratories, France). Grafts for *ex-vivo* mechanical stress quantification were performed in NSG mice from our own breeding. Mice were housed and bred under specific pathogen-free (SPF) conditions in an accredited animal facility. Animals were maintained on a 12-hour light/dark cycle (lights on at 07:00) at a controlled temperature of 22 ± 1 °C and relative humidity of 55 ± 5%. Food and water were provided ad libitum. Cages were enriched with wood shavings, bedding, and at least one additional item (cardboard structure, igloo, or sizzle nest tube) to promote environmental enrichment.

### Statistical analysis

Comparisons between two experimental groups were performed using a paired two-tailed Student t-test (*in vitro*) and Mann-Whitney tests (*in vivo*). Comparisons between more than two experimental groups were performed using two-ways ANOVA. p-value<0.05 was considered significant. All analysis was performed using GraphPadPrism 10.1.2 software.

## Acknowledgments

We thank our colleagues for their critical reading of the manuscript. We thank the Pôle technologique du Centre de Recherche en Cancérologie de Toulouse (Manon Farcé, Laetitia Ligat, Laura Lestienne, Tiphaine Fraineau, Frédéric Pont, Emeline Sarot), for the technical support in imaging, vectorology, proteomics and cytometry platforms. We thank the I2MC We-Met platform in Toulouse (Alexandre Lucas and Corinne Bernis) for their help in simple western experiment and analysis. We thank Christèle Segura from I2MC histology platform. We thank Rémi Coursson from LAAS-CNRS for his support in 3D printing. We acknowledge Yvan Nicaise for providing bioinformatics expertise and help for histology. We address a special thanks to Cedric Baudelin, Emilie Sinhlivong, Tristan Chapon, Amanda Corini, Pierre-Paul Colombo and Ibou Ka from animal facilities, CREFRE-Toulouse (Centre régional d’exploration fonctionnelle et de ressources expérimentales), as well as ENI (Carine Pestouri) for their help and support for all in vivo experiments and non-invasive imaging. Views and opinions expressed are, however, those of the author only and do not necessarily reflect those of the European Union or the European Research Council. Neither the European Union nor the granting authority can be held responsible for them.

## Fundings

Fondation Toulouse Cancer Santé (Mecharesist)

Inserm Plan Cancer (PressDiagTherapy; MECHAEVO; MEchanoStem)

Inca PLBIO INCa_16095

Toucan ANR Laboratory of Excellence

MSCA-ITN/ETN PIPgen (Project ID: 955534)

Fondation ARC (ARCPJA2021060003932, ARCPGA2022120005630_6362-3)

Université de Toulouse Emergence (MECACAN) Cancéropôle GSO Emergence de Consortium

ERC Starting grant UnderPressure grant agreement number 101039998

## Author contributions

Experimentation, analysis, visualisation and interpretation of data: MDL, SA, NT, TM, RDA, BT, PS, PA, MD and JGG.

Supervision of students: MDL, PA, JGG Supervision of staff: JGG

Methodology: MDL, SA, NT, TM, RDA, BT, PS, PA, MD and JGG

Drafting of the manuscript: MDL and JGG Revising the manuscript: all authors

Conceptualization of funded project, obtained funding, project administration, project supervision, validation: MDL, PA, MD and JGG

## Data and materials availability

All data are available in the main text or supplementary materials.

We generated and stored bulk RNA sequencing transcriptomic data available upon request.

## Supplementary Materials

**Movie S1. 3D visualization of compressive device.** Blender 4.5 software was used for 3D visualization.

