## Supplemental Figures for "The tumor suppressor p53 mutational status controls epithelial 3D cell growth under mechanical compression"

Supp figure 1

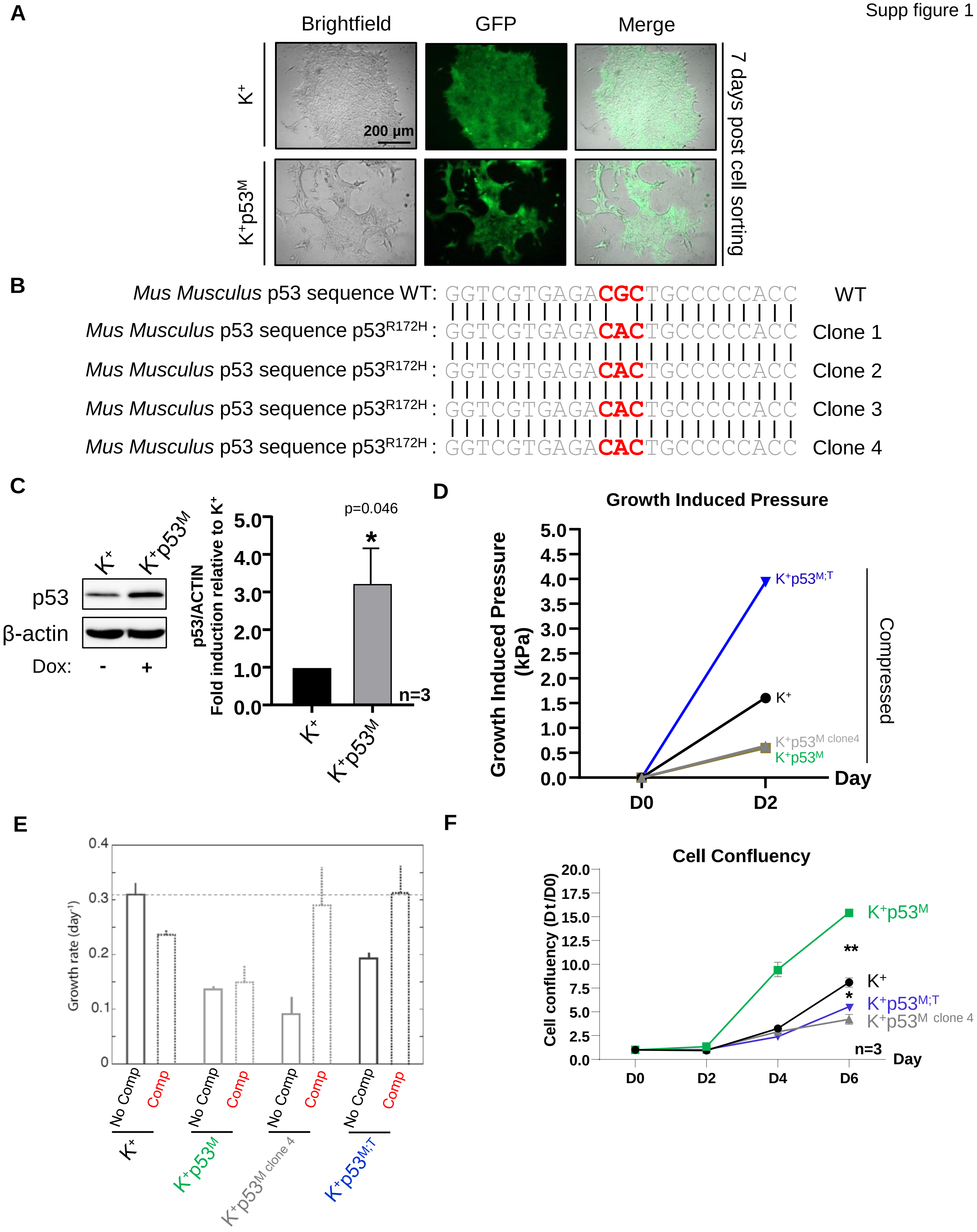

Supp figure 2

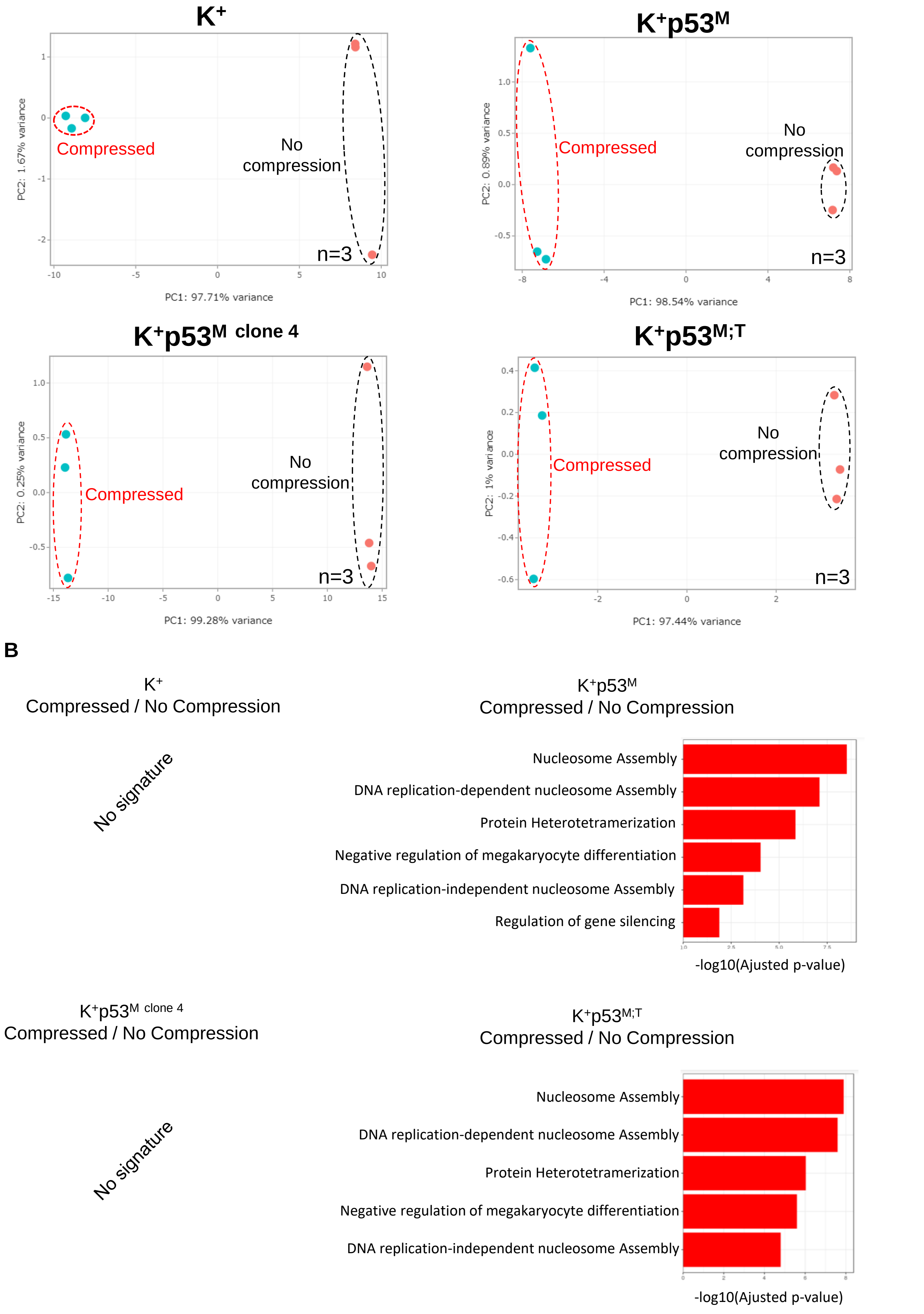

A

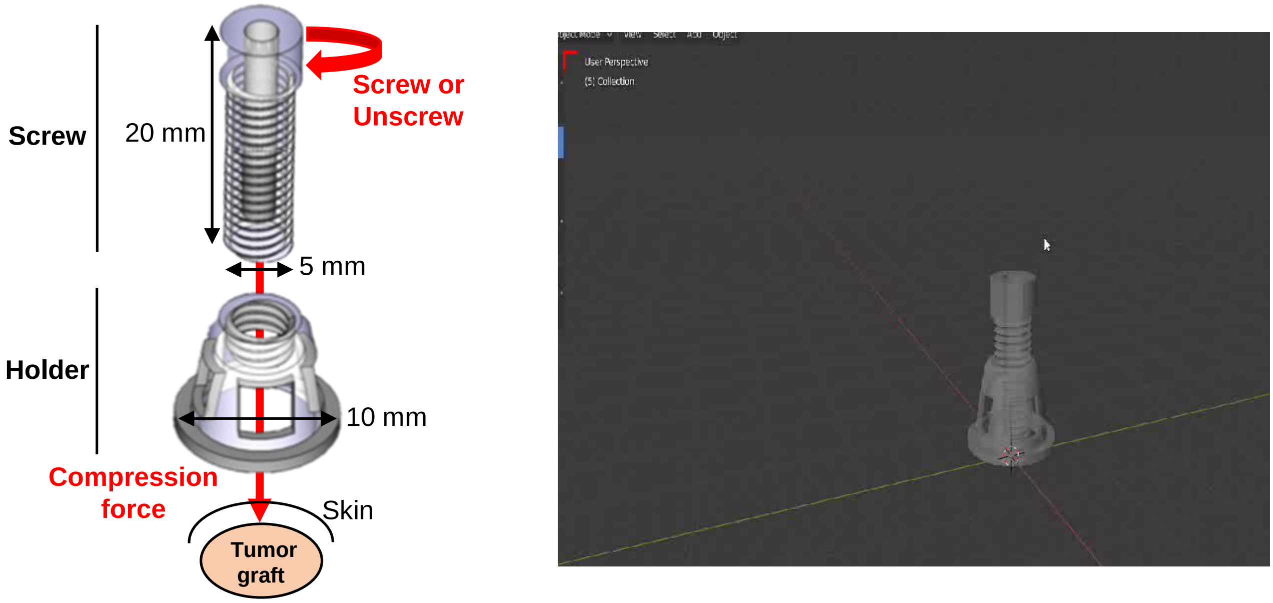
Screw
20 mm
Screw or
Unscrew
5 mm
Holder
10 mm
Compression
force
Skin
Tumor
graft

B

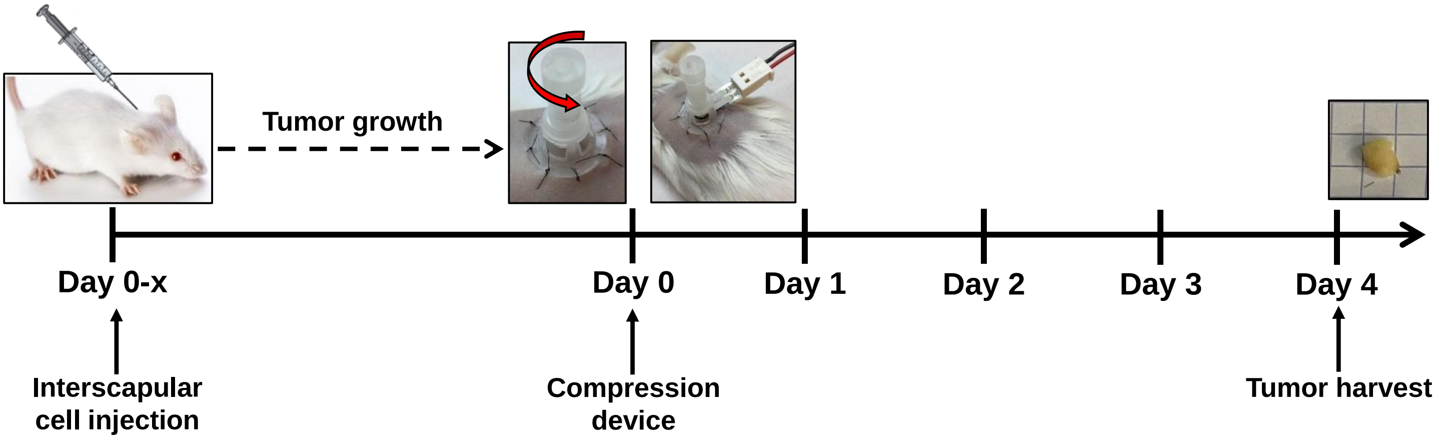
Tumor growth
Day 0-x
Day 0
Day 1
Day 2
Day 3
Day 4
Interscapular
cell injection
Compression
device
Tumor harvest

### Tumor detection with indicated genotype

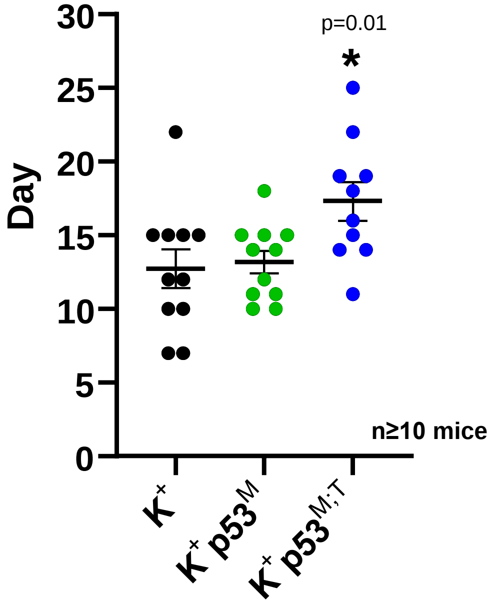

### *c-Fos* and *FosB* mRNA expression

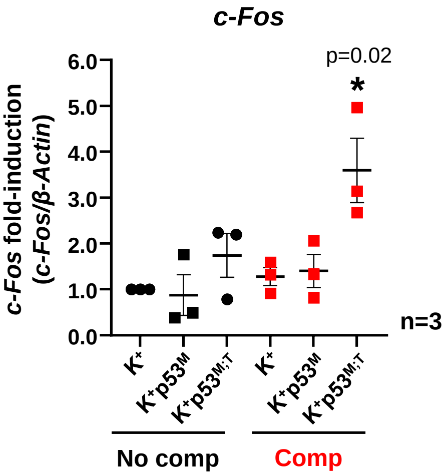


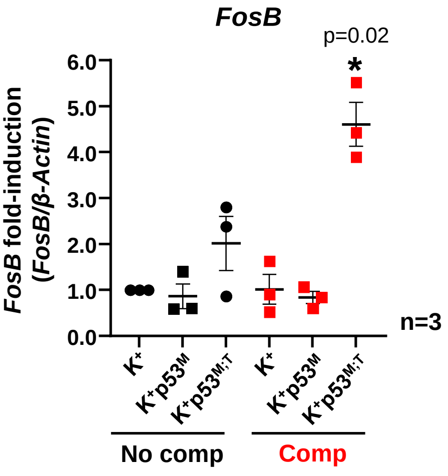

**A**

### YAP pathway

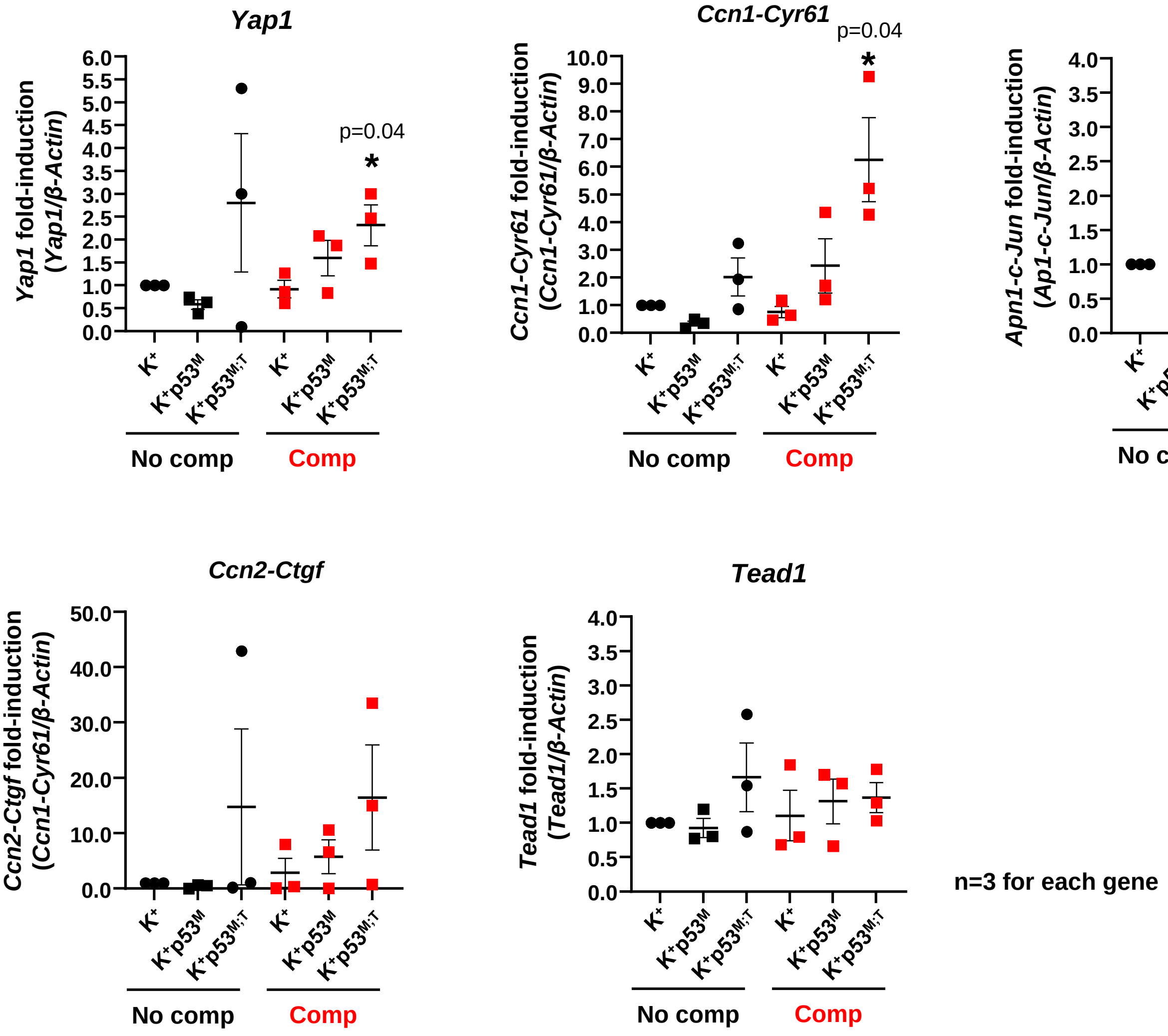


n=3 for each gene

**B**

### Hypoxia

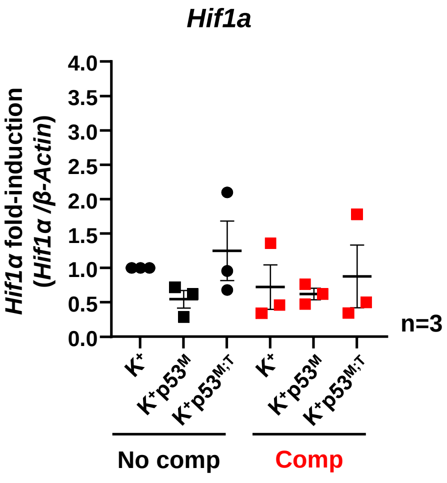


n=3
