## Supplemental Tables for "The tumor suppressor p53 mutational status controls epithelial 3D cell growth under mechanical compression"

### Supporting Tables

#### *Supporting Table 1. List of genes overexpressed in $K^{+}$ $p53^{M;T}$ spheroids under confinement*

| Symbol | Fold change $K^{+}$ Comp/No comp | p-value adj | Fold change $K^{+}p53^{M}$ Comp/No comp | p-value adj | Fold change $K^{+}p53^{M;T}$ Comp/No comp | p-value adj |
|---|---|---|---|---|---|---|
| ***Fos*** | **3.214** | **1.28E-47** | **3.219** | **2.43E-62** | **8.212** | **3.59E-214** |
| *Sfrp2* | NA | NA | NA | NA | 7.877 | 7.08E-40 |
| *P2ry1* | 1.135 | 2.85E-01 | NA | NA | 5.328 | 1.27E-04 |
| *Acss3* | 5.306 | 4.98E-22 | 59.987 | 2.14E-40 | 4.840 | 1.41E-04 |
| *Rny3* | 4.352 | 4.75E-05 | 5.476 | 2.72E-08 | 4.673 | 6.88E-05 |
| ***Fosb*** | **0.823** | **1.26E-03** | **1.27** | **6.91E-06** | **4.531** | **0.00E+00** |
| *Apol8* | 0.101 | 8.86E-10 | NA | NA | 4.506 | 1.45E-03 |
| *Adgrl2* | 220.169 | 8.52E-10 | NA | NA | 3.966 | 5.94E-06 |
| *Fam180a* | NA | NA | NA | NA | 3.298 | 1.60E-17 |
| *Hist1h4a* | 4.077 | 1.01E-06 | 8.887 | 3.66E-23 | 3.272 | 6.26E-08 |
| *Rmrp* | 10.029 | 4.89E-76 | 8.396 | 2.05E-187 | 3.258 | 3.67E-37 |
| *Hist1h4b* | 3.775 | 1.58E-05 | 4.461 | 6.36E-10 | 3.216 | 6.07E-06 |
| *Gpr25* | NA | NA | NA | NA | 3.138 | 3.08E-05 |
| *Hist1h4c* | NA | NA | 9.322 | 3.49E-09 | 3.127 | 3.97E-04 |
| *Wscd2* | NA | NA | NA | NA | 3.046 | 1.51E-02 |

#### *Supporting Table 2. List of primers and siRNA*

| APPLICATION | GENE | SEQUENCE |
|---|---|---|
| Sub-cloning | *Mus Musculus* | |
| | *TP53 (53kDa)*<br>*SfiI restriction enzyme* | Forward:<br>5'-AAAGGCCTCTGAGGCCATGACTGCCATGGAGGAGTC-3'<br>Reverse:<br>5’- AAAGGCCTGACAGGCCTCAGTCTGAGTCAGGCCC-3' |
| | $TP53^{R172H ;R210*}$<br>($p53^{M,T}$) | Forward:<br>5'-AAAGGCCTCTGAGGCCATGACTGCCATGGAGGAGTC-3'<br>Reverse:<br>5’-AAGGCCTGACAGGCCTCAAAAAGTCTGCCTGTCTTCCAG-3' |
| Directed Mutagenesis | $p53^{R172H}$<br>($p53^{M}$) | Forward: 5'-GAGGTCGTGAGACGCTGCCCCCACCATG-3'<br>Reverse: 5’-CATGGTGGGGGCAGCGTCTCACGACCTC-3' |
| Sequencing | *pSB-tet-GN* | Forward: 5'-GCAGAGCTCGTTTAGTGAAC-3'<br>Reverse: 5’-GCAATAGCATCACAAATTTCAC-3' |
| RT-qPCR | *Pik3ca* | Forward: 5’-CTGCAGTTCAACAGCCACAC-3'<br>Reverse: 5’-TCTCGCCCTTGTTCTTGTCC-3' |
| | *Pik3cb* | Forward: 5’-CTGATTTTACGGCGGCATGG-3'<br>Reverse: 5’-TGAGGGCCTCGTCAAACTTC-3' |
| | *Pik3cd* | Forward: 5'-GTCCACTCCTCCTCCATCCT-3'<br>Reverse: 5’-GAGGTTTGGCACGTGGTTTC-3' |
| | *Pik3cg* | Forward: 5'-CGACCGAAAGTTCAGGGTCA-3'<br>Reverse: 5’-CAATAGAGCCCCTTTGGGCA-3' |
| | *Ap1-c-Jun* | Forward: 5'-CCGGACTGTTCATCCGTTTG-3'<br>Reverse: 5’-CCCGGACTTGTGAGCTTC-3' |
| | *Actin* | Forward: 5’-CAGCCTTCCTTCTTGGGTATG-3'<br>Reverse: 5’-GGCATAGAGGTCTTTACGGATG-3' |
| | *c-Fos* | Forward: 5'-GTGAAGACCGTGTCAGGAG-3'<br>Reverse: 5’-GTGTATCTGTCAGCTCCCTC-3' |
| | *Ccn1-Cyr61* | Forward: 5'-CAATTTCGGCGCCAGCTC-3'<br>Reverse: 5’-GACAGCCCAGATTGGGGAG-3' |
| | *Ccn2-Ctgf* | Forward: 5'-CAACCGCAAGATCGGAGTG-3'<br>Reverse: 5’-GGCAGCTTGACCCTTCTC-3' |
| | *Fosb* | Forward: 5'-CTTCAACCAGCACAACCACC-3'<br>Reverse: 5’-GAAGTCGATCTGTCAGCTCCC-3' |
| | *Hif1α* | Forward: 5'-GTGTGAGAAAACTTCTGGATGC-3'<br>Reverse: 5’-CCCCATGTATTTGTTCACGTTATC-3' |
| | *Tead1* | Forward: 5'-CTGGCTATCTATCCGCCGTG-3'<br>Reverse: 5’-CTTCCTGGTCCTTGTCTTTCCC-3' |
| | *Yap1* | Forward: 5'-GATGGCCAAGACATCTTCTGGTC-3'<br>Reverse: 5’-GAATTCATCAGCGTCTGGGG-3' |
| SiRNA | | |
| | *SiRNA-Scramble*:<br>*ON-TARGET plus Non-targeting Pool*<br>*(D-001810-10-20)* | UGGUUUACAUGUCGACUAA<br>UGGUUUACAUGUUGUGUGA<br>UGGUUUACAUGUUUUCUGA<br>UGGUUUACAUGUUUUCCUA |
| | *SiRNA-c-Fos*:<br>*ON-TARGET plus Mouse Fos (14281) siRNA SMART Pool*<br>*(L-041157-00-0010)* | GCGCAGAGCAUCGGAGAA<br>GGAGGAGGAGCUGACAGA<br>GGAUUUGACUGGAGGUCUG<br>GCGCAGAUCUGUCCGUCUC |
| | *SiRNA-Fosb*:<br>*ON-TARGET plus Mouse Fos (14282) siRNA SMART Pool*<br>*(L-045464-01-0010)* | GUGAAUGAGUGGUCGGAUU<br>CCGCUAAGGAAGACGGCUU<br>GUUGUUAGCCCUUCGUACA<br>CCAUCUUGCUGGAGCGCUU |

#### *Supporting Table 3. List of primary antibodies and reagents*

| PRIMARY ANTIBODY | SPECIES | SOURCE | REFERENCE NUMBER | DILUTION WB | DILUTION IHC/IF |
|---|---|---|---|---|---|
| β-ACTIN | Mouse | Merck | #A2228 | 1/10000 | 1/1000 |
| AKT | Rabbit | Cell Signaling | #4691 | 1/1000 | |
| Cleaved-Caspase 3 | Rabbit | Cell Signaling | #9664 | | 1/100 (IHC 1/600(IF) |
| c-FOS | Rabbit | Cell Signaling | #2250 | 1/1000 | |
| DAPI | | Merck | #D9542 | | 1/1000 |
| FOSB | Rabbit | Cell Signaling | #2251 | 1/1000 | |
| GFP | Goat | Abcam | #ab6673 | | 1/250 |
| Ki67 (IHC) | Rabbit | Abcam | ab15580 | | 1/100 |
| Ki67(IF) | Rabbit | Thermofisher | MA5-14520 | | 1/500 |
| p53 | Rabbit | Cell Signaling | #2524 | 1/1000 | 1/1000 |
| p53 | Rabbit | Abcam | #131442 | 1/1000 | |
| p-AKT(Ser473) | Rabbit | Cell Signaling | #4060 | 1/2000 | |
| YAP | Mouse | Santa Cruz | #101199 | | 1/400(IF) |

#### *Supporting Table 4. List of secondary antibodies and reagents*

| SECONDARY ANTIBODY | SPECIES | SOURCE | REFERENCE NUMBER | DILUTION WB | DILUTION IHC/IF |
|---|---|---|---|---|---|
| Anti-rabbit IgG, HRP-linked Antidoby | Goat | Cell Signaling | #7074 | 1/7500 | |
| Anti-mouse IgG-Horse Radish Peroxidase | Goat | Invitrogen | #31430 | 1/7500 | |
| Anti-rabbit Alexa Fluor 488 | Goat | Abcam | #ab150077 | | 1/1000 |
| Anti-rabbit Alexa Fluor 568 | Goat | Abcam | #ab175470 | | 1/1000 |
| ImmPRESS® HRP Horse Anti-Rabbit IgG Polymer Detection Kit, Peroxidase | Horse | Vector lab | #MP-7401 | | |
